# BAF complex-independent gene activation by SS18::SSX

**DOI:** 10.64898/2026.01.26.701739

**Authors:** Afroditi Sotiriou, Jinxiu Li, Sanya Middha, Jake A. Ward, Selina Troester, Lianghao Mao, Martin Schneider, Dario Frey, Eliza C. Wray, Sara Bocedi, Kyllie Smith-Fry, Linda Morrison, Lara Carroll, Mihaly Badonyi, Joseph A. Marsh, Ashok Kumar Jayavelu, Cristina Mayor-Ruiz, Bradley R. Cairns, Kevin B. Jones, Nezha S. Benabdallah, Ana Banito

**Author notes:** Correspondence to: N.S.B: Institute of Genetics and Cancer, University of Edinburgh, Western General Hospital, Crew Road South, EH4 2XR, Edinburgh, United Kingdom.; A.B: German Cancer Research Center (DKFZ), 69120 Heidelberg, Germany.

## Abstract

In synovial sarcoma, the BAF subunit SS18 is fused to SSX, a transcriptional repressor, generating the oncogenic SS18::SSX fusion protein. Incorporation of SS18::SSX into BAF complexes leads to their aberrant retargeting to Polycomb-repressed genes via SSX, while simultaneously altering their composition and activity. The presence of BAF at Polycomb target sites is widely assumed to be essential for gene activation. Here, we directly tested the requirement for BAF activity in synovial sarcoma cell survival and SS18::SSX-driven transcription. Using targeted degradation of BAF ATPase subunits and deletion of core components, we show that BAF loss has modest effects on sarcoma cell viability and does not impede SS18::SSX target gene expression. Consistently, deletion of the BAF ATPase subunit *Smarca4* does not impair SS18::SSX-driven tumor formation *in vivo*. Using domain-specific SS18::SSX mutants, we further demonstrate that the fusion can activate oncogenic transcription independently of BAF interaction, and that this activity depends on the C-terminal QPGY-rich domain of SS18. Mechanistically, SS18::SSX promotes transcription by engaging the histone acetyltransferase EP300, independently of BAF. Accordingly, pharmacologic degradation of EP300/CREBBP suppresses SS18::SSX-driven transcriptional programs and impairs synovial sarcoma cell survival. Together, these findings challenge the view that BAF activity is required for SS18::SSX-mediated transcriptional activation and demonstrate that aberrant Polycomb target gene expression is sustained through recruitment of transcriptional coactivators in the absence of BAF. Our work reveals new therapeutic vulnerabilities in synovial sarcoma and suggests broader relevance for targeting coactivator-dependent transcription in fusion-driven cancers.

**Highlights:**

- BAF degradation does not alter SS18::SSX-activated transcriptional programs
- Direct SS18::SSX transcriptional activation is independent of BAF interaction
- The SS18 C-terminus engages the co-activator EP300 to promote gene expression
- Small molecule degraders of EP300/CREBBP abolish SS18::SSX-mediated transcription

## INTRODUCTION

Synovial sarcoma represents a paradigm of BAF-dysregulated cancer. BAF (or mSWI/SNF) chromatin-remodeling complex is a central ATP-dependent regulator of genome organization that plays essential roles in establishing and maintaining lineage-specific transcriptional programs^1-4^. The catalytic ATPase subunits SMARCA2 (BRM) and SMARCA4 (BRG1) reposition or eject nucleosomes to regulate enhancer and promoter accessibility^3,5^.

In synovial sarcoma, a recurrent chromosomal translocation between chromosome 18 and chromosome X generates the oncogenic SS18::SSX fusion protein, which is sufficient to drive tumorigenesis^6-12^. Early studies established that SS18 is a core component of the BAF chromatin-remodeling complex and that SS18::SSX bridges BAF to Polycomb-occupied loci through the high affinity of the SSX C-terminal RD domain to H2AK119ub1-rich regions deposited by the non-canonical Polycomb Repressive Complex, ncPRC1.1^13-16^. SS18::SSX stably incorporates into BAF, displacing and promoting degradation of wild-type SS18 and destabilizing canonical BAF (cBAF) complexes, resulting in increased representation of fusion-containing non-canonical GLTSCR1-containing BAF (GBAF, or ncBAF) complexes^17-20^. Together, these findings led to a model in which SS18::SSX drives synovial sarcoma pathogenesis through both aberrant targeting of BAF to normally repressed chromatin and altered BAF complex composition, resulting in cBAF depletion. Recent work has highlighted BRD9, a defining subunit of the ncBAF complex, as synthetically lethal in synovial sarcoma^21,22^. However, BRD9 removal did not reduce SS18::SSX-activated transcription^22^. Efforts focused on BRD9 degraders, such as FHD-609 (NCT04965753) and CFT8634 (NCT05355753) in patients with advanced synovial sarcoma, however, their impact in clinical trials was modest and produced unexpected toxicities^23,24^. Despite that many preclinical studies in so-called BAF-addicted cancers (i.e. uveal melanoma and SMARCA4-mutated non-small cell lung cancer) using SMARCA2/SMARCA4 degraders and inhibitors have demonstrated that acute BAF ATPase blockade induces profound transcriptional and chromatin alterations and showed promising results in clinical trials^25-28^, this therapeutic strategy has not been applied yet to synovial sarcoma.

Here, we sought to investigate the requirement of the BAF complex in this disease both at the levels of SS18::SSX-activated transcription and cell viability. We demonstrate that pharmacologic or genetic perturbation of BAF complexes mildly affects synovial sarcoma cell viability and does not abrogate SS18::SSX mediated transcriptional programs. Instead, SS18::SSX acts as a chimeric transcription factor by engaging EP300 in the absence of BAF interaction. Accordingly, SS18::SSX-driven transcription is efficiently suppressed by the dCBP-1 degrader and results in enhanced sensitivity of synovial sarcoma cells to dCBP-1 treatment.

Our findings challenge the long-standing notion that BAF is critical for SS18::SSX transcriptional activation of Polycomb target genes and position EP300/CREBBP as a critical coactivator that sustains SS18::SSX-driven transcription, offering a promising avenue for targeted therapeutic intervention.

## RESULTS

### Synovial sarcoma cell lines exhibit limited reliance on BAF complex enzymatic activity

In the current model of SS18::SSX-driven transcriptional dysregulation, the SS18 moiety mediates incorporation into the BAF complex, whereas the SSX region directs recognition of PRC1 target regions marked by H2AK119ub1^13-16^. This mistargeting of BAF complexes is thought to result in activation of Polycomb target genes. To investigate the extent to which BAF complex enzymatic activity contributes to SS18::SSX function, we used the small-molecule degrader ACBI1, which potently degrades the ATPase subunits SMARCA2/SMARCA4, rendering the complex enzymatically inactive^29^. Treatment of HSSY-II synovial sarcoma cells with 500 nM ACBI1 for 72 h resulted in efficient degradation of SMARCA2 and SMARCA4 (**Figure 1A**). Immunoprecipitation (IP)-mass spectrometry (MS) of the endogenous oncofusion following ACBI1 treatment revealed that SMARCA2/4 degradation was accompanied by loss of interaction with multiple BAF subunits, consistent with previous work showing that ACBI1 can induce complex disassembly^29^ (**Figure 1B, 1C**).

**Figure 1:**
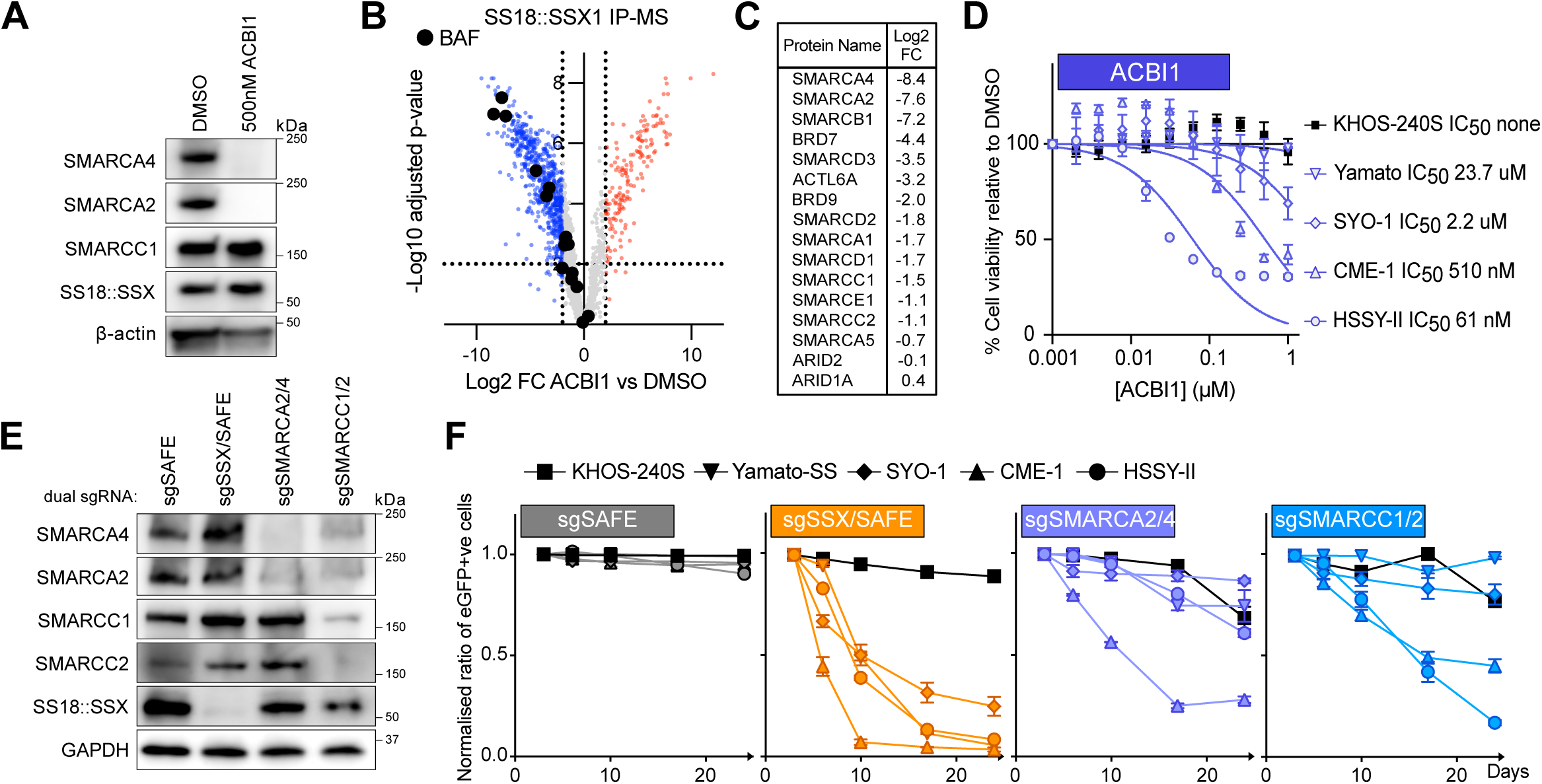
BAF removal has a moderate effect in synovial sarcoma cell line survival. **(A)** Western Blot of HSSY-II whole cell extracts treated for 72h with DMSO or 500nM ACBI1 revealed using SMARCA4, SMARCA2, SMARCC1, SS18::SSX or β-actin antibodies. n = 1. **(B)** Volcano plots of mass spectrometry data following SS18::SSX pull down in HSSY-II synovial sarcoma cells treated for 72h with DMSO or 500nM ACBI1, normalised to interactome of DMSO treated cells. Data represents log2 fold change enrichment plotted on the x-axis and -log10 (adjusted p value) plotted on the y-axis. Blue/Red dots indicate lost/gained interactors upon ACBI1 treatment. Black dots indicate BAF members. n = 4 biological replicates. **(C)** List of BAF members present in interactome dataset and their associated log2 fold change value in ACBI1 SS18::SSX pull downs normalised to DMSO. Listed members are shown as black dots in (B). **(D)** Viability of 4 synovial sarcoma cell lines (Yamato-SS, SYO-1, CME-1, HSSY-II) and an osteosarcoma cell line (fusion negative control) KHOS-240S treated with various concentrations of ACBI1. The symbols represent the mean ± S.E.M of n = 3 biological replicates, the solid line represent the nonlinear curve fitting and its associated IC50. **(E)** Western Blot of HSSY-II-Cas9 whole cell extracts expressing a safe sgRNA as control or with double guides targeting SSX + a control region (sgSSX/SAFE), SMARCA2 + SMARCA4 (sgSMARCA2/4) or SMARCC1 + SMARCC2 (sgSMARCC1/2) for 7 days revealed using SMARCA4, SMARCA2, SMARCC1, SMARCC2, SS18::SSX or GAPDH antibodies. Representative of n = 2 biological replicates. **(F)** Cell competition assay performed in the osteosarcoma cell line KHOS-240S-Cas9 or in the 4 synovial sarcoma lines expressing Cas9 expressing a safe sgRNA as control or double guides targeting SSX + a control region (sgSSX/SAFE), SMARCA2 + SMARCA4 (sgSMARCA2/4) or SMARCC1 + SMARCC2 (sgSMARCC1/2). Data represents the mean ± S.E.M of n = 2 biological replicates.

We previously showed that ACBI1 treatment does not disrupt SS18::SSX distribution to chromatin sites marked by Polycomb^16^. We therefore set out to examine how removing BAF enzymatic activity from these sites (and generally) affects cell viability. We tested the effect of 5 days ACBI1 treatment on a panel of synovial sarcoma cell lines (HSSY-II, SYO-1, CME-1 and Yamato-SS) as well as an osteosarcoma cell line, KHOS-240S, as control. We observed a stronger dependency of synovial sarcoma cell lines compared to the control cell line KHOS-240S, but treatment with ACBI1 resulted in only a moderate reduction in viability of the synovial sarcoma lines and relatively high IC50 values (**Figure 1D**). Orthogonally, we used genetic double knockouts of *SMARCA2* and *SMARCA4* by CRISPR/Cas9 (sgSMARCA2/4) and of *SMARCC1* and *SMARCC2* (sgSMARCC1/2), which are core BAF subunits required for proper complex assembly on chromatin^5^. As controls, we used either a non-targeting sgRNA (sgSAFE) or knockout of the fusion (sgSSX). Of note, disruption of *SMARCC1* and *SMARCC2* also led to depletion of SMARCA2 and SMARCA4 at the protein level, likely by destabilizing the complex (**Figure 1E**). Again, sgSMARCA2/4 or sgSMARCC1/2 caused only mild or no proliferative disadvantage, despite efficient CRISPR editing (**Figure 1F, Figure S1A,B**). In contrast, sgSSX markedly led to reduction of cell proliferation over time as expected. This demonstrates that ablation of BAF complexes does not phenocopy loss of SS18::SSX, suggesting instead that the fusion can act independently of the BAF complex.

### BAF complex disruption does not suppress SS18::SSX-mediated transcriptional program

We then examined how BAF perturbation affects gene expression by performing RNA-sequencing in the synovial sarcoma cell line HSSY-II. To our surprise, while sgSSX elicited strong global transcriptional changes, neither pharmacologic nor genetic disruption of BAF caused major changes in gene expression (**Figure 2A**). A closer look at classic SSX target genes, mainly Polycomb targets, confirmed that their expression was reduced by oncofusion depletion, but not by BAF removal (**Figure 2B**). To further assess this across the top 500 genes downregulated by SS18::SSX loss, we performed Gene Set Enrichment Analysis (GSEA) and confirmed that BAF removal did not produce concordant or statistically significant changes in gene expression (**Figure S2A**). This discrepancy between the effects of oncofusion removal and BAF depletion was similarly observed in other synovial sarcoma cell lines, including Yamato-SS (**Figure 2C**), CME-1, and SYO-1 (**Figure S2B-D**).

**Figure 2:**
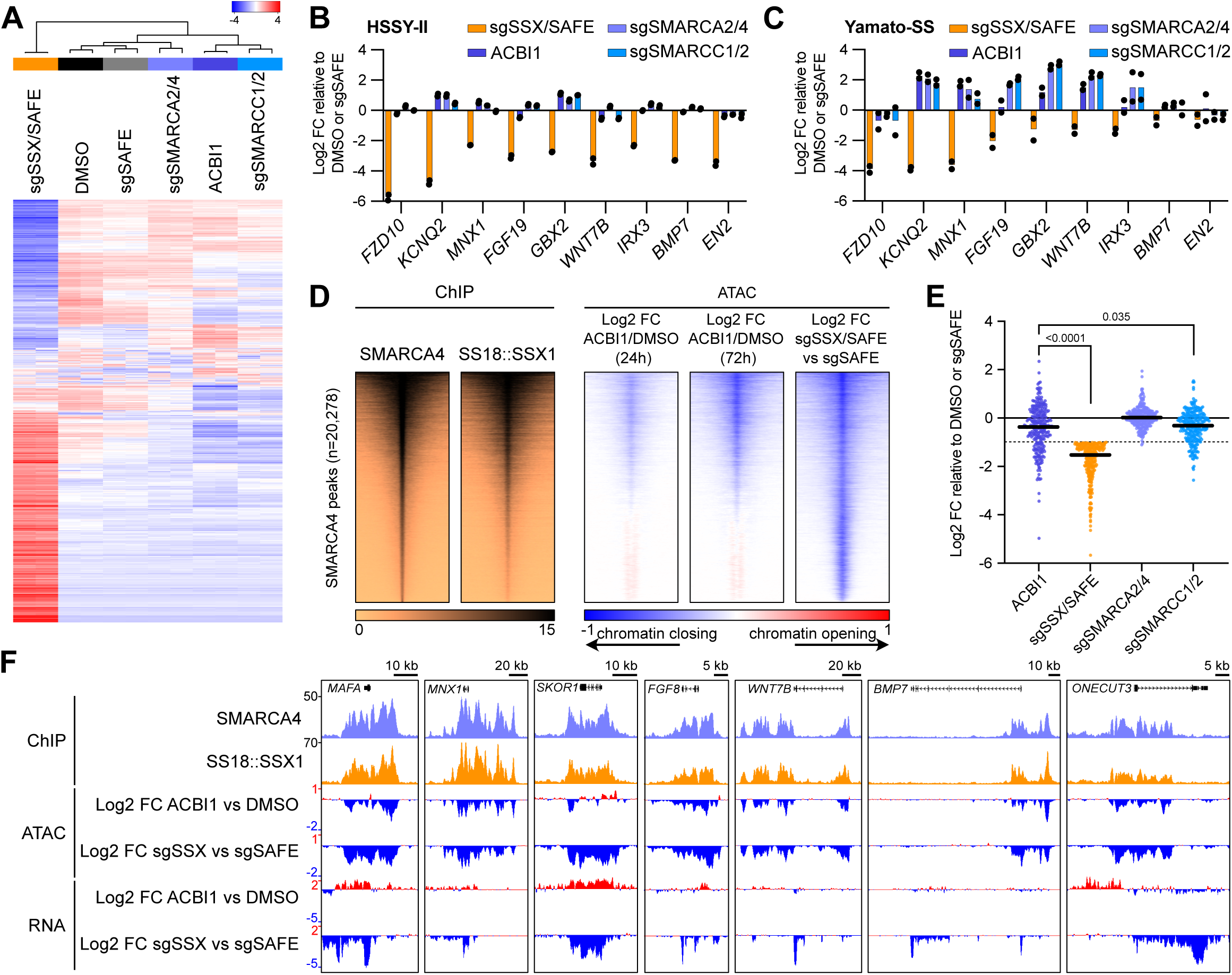
SS18::SSX-dependent gene expression persists despite BAF complex disruption. **(A)** RNA-seq heatmap showing the 1,000 most variable genes with a cutoff z-score of 4 from HSSY-II cells treated for 72h with DMSO or 500nM ACBI1 or of HSSY-II-Cas9 cells expressing a safe sgRNA as control or with double guides targeting SSX + a control region (sgSSX/SAFE), SMARCA2 + SMARCA4 (sgSMARCA2/4) or SMARCC1 + SMARCC2 (sgSMARCC1/2). n=2 biological replicates. **(B)** Log2-transformed fold change of FPKM values in HSSY-II RNA-seq relative to DMSO or sgSAFE over selected SS18::SSX target genes. Data represents the mean. **(C)** qRT-PCR displaying log2-transformed fold change of mRNA levels normalised by GAPDH in Yamato-SS cells treated for 72h with DMSO or 500nM ACBI1 or of Yamato-SS Cas9 cells expressing sgSAFE, (sgSSX/SAFE), (sgSMARCA2/4) or (sgSMARCC1/2) relative to DMSO or sgSAFE for 7 days. Data represents the mean of n=2 biological replicates. **(D)** Left: heatmaps for SMARCA4 and SS18::SSX1 (endogenously HA tagged) ChIP-seq from Banito et al., 2018 over SMARCA4 peaks (n = 20,278). Rows correspond to ±5-kb regions across the midpoint of each SMARCA4-enriched region, ranked by increasing signal. Right: ATAC-seq heatmaps displaying log2-transformed fold change of ACBI1 treated cells over DMSO after 24h or 72h of treatment and of sgSSX/SAFE expressing cells over sgSAFE cells. n=2 biological replicates. **(E)** Scatter plot of log2-transformed fold change of FPKM values in HSSY-II RNA-seq relative to DMSO or sgSAFE over genes associated with SMARCA4 occupancy and showing <-1.5 fold down-regulation upon sgSSX/SAFE knockout. Biological replicates are combined using their mean. Bold line represents median and dotted line represents -1.5 cutoff. p-values represent two-tailed Wilcoxon test. **(F)** Gene tracks for SMARCA4, SS18::SSX1 ChIP-seq, Log2-Fold changes of ATAC-seq or Log2-Fold changes of RNA-seq at loci from (E) such as *MAFA*, *MNX1*, *SKOR1*, *FGF8*, *WNT7B*, *BMP7* and *ONECUT3*.

We next determined the genome-wide SMARCA4 chromatin occupancy intersecting with SS18::SSX1 distribution by ChIP-seq, to identify BAF occupied sites for further interrogation (**Figure 2D**). BAF removal affects chromatin accessibility and leads to a change in chromatin opening by ATAC-seq after ACBI1 treatment for either 24 or 72 hours (h) in HSSY-II synovial sarcoma cells. ACBI1 treatment predominantly altered chromatin accessibility at peaks coincident with high BAF and high SS18::SSX occupancy. Yet, the magnitude of these changes was substantially lower than the chromatin closing observed upon SS18::SSX removal (**Figure 2D**). At RefSeq gene promoters, we observed no closing and even some increase in chromatin opening upon ACBI1 treatment. In contrast, sgSSX led to strong chromatin closing at promoters occupied by SS18::SSX and SMARCA4 (**Figure S2E**). Next, we investigated how regions co-occupied by SMARCA4 and SS18::SSX relate to changes in gene expression. To this end, we assigned SMARCA4 peaks to putative genes using GREAT analysis. Focusing on the genes downregulated upon SS18::SSX removal, we compared log2 fold changes following either ACBI1 treatment, or knockout of BAF components relative to the DMSO or sgSAFE controls. Indeed, neither ACBI1 treatment nor genetic disruption of BAF components resulted in gene expression changes comparable in magnitude to those observed upon fusion knockout (**Figure 2E**). Together our data show that BAF complex disruption, even when accompanied by partial chromatin closing, does not substantially affect expression of oncofusion target genes (**Figure 2F**).

### SS18::SSX can induce transcription and chromatin opening independently of BAF catalytic activity

Because synovial sarcoma cells display inconsistent sensitivity to BAF removal, without substantial alterations in SS18::SSX-driven expression signatures, we considered if this might reflect a cell-type-specific result, rather than a direct molecular requirement of the fusion protein itself. To explore this further, we ectopically expressed SS18::SSX1 in the osteosarcoma cell line KHOS-240S, which lacks the fusion and is insensitive to both BAF (Figure 1D, 1F) and Polycomb complex depletion^16^. To probe initiation of the fusion’s transcriptional program in the absence of BAF, we pre-treated KHOS-240S cells with ACBI1 to deplete BAF prior to introducing the fusion, thereby directly testing whether BAF is required for the establishment of SS18::SSX-driven transcription (**Figure 3A**).

**Figure 3:**
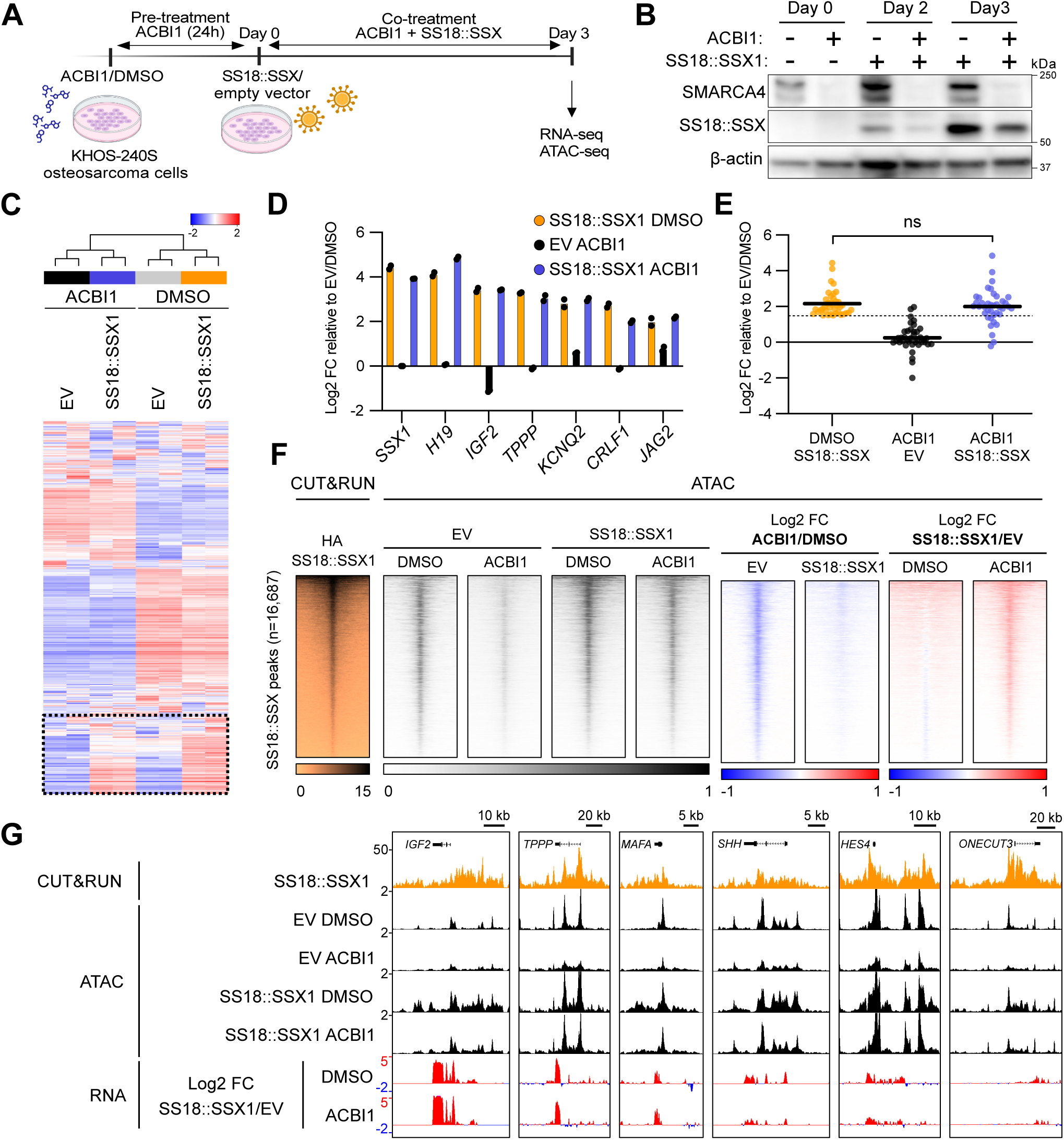
SS18::SSX activates transcription and opens chromatin independently of BAF. **(A)** Schematics showing timeline for ACBI1/DMSO treatment and SS18::SSX/EV delivery in KHOS-240S cells. **(B)** Western Blot of KHOS-240S whole cell extracts after 24h pre-treatment with DMSO or 500nM ACBI1 (Day 0), or at day 2 and 3 following expression of an empty vector (EV) or SS18::SSX1 transgene and continued treatment with DMSO or ACBI1. **(C)** RNA-seq heatmap showing the 1,000 most variable genes with a cutoff z-score of 2 from KHOS-240S cells pre-treated with DMSO or 500nM ACBI1 for 24h prior to expression of an empty vector (EV) or SS18::SSX1 transgene for additional 72h and co-treatment with DMSO or ACBI1. n=2 biological replicates. Dotted square highlights genes up- regulated by SS18::SSX1 expression **(D)** Log2-transformed fold change of FPKM values in KHOS-240S RNA-seq relative to EV/DMSO over selected SS18::SSX target genes. Data represents the mean. **(E)** Scatter plot of log2-transformed fold change of FPKM values in KHOS-240S RNA-seq relative to EV/DMSO over genes up-regulated by SS18::SSX1 expression and showing >1.5 fold up-regulation. Biological replicates are combined using their mean. Bold line represents median and dotted line represents 1.5 cutoff. p-values represent two-tailed Wilcoxon test. **(F)** Left: heatmaps for SS18::SSX1 (HA tagged-transgene) CUT&RUN from Benabdallah et al., 2023 over SS18::SSX1 peaks (n = 16,687). Rows correspond to ±5-kb regions across the midpoint of each SS18::SSX1-enriched region, ranked by increasing signal. Right: ATAC-seq heatmaps (in greys) of KHOS-240S cells pre-treated with DMSO or 500nM ACBI1 for 24h prior to expression of an empty vector (EV) or SS18::SSX1 transgene for additional 72h and co-treatment with DMSO or ACBI1 or ATAC-seq heatmaps (in blue-red gradient) of comparison using log2- transformed fold change of either ACBI1 relative to DMSO in EV or SS18::SSX1 condition or SS18::SSX relative to EV in DMSO or ACBI1 conditions. n-2 biological replicates. **(G)** Gene tracks for SS18::SSX1 CUT&RUN, ATAC-seq of KHOS-240S cells pre-treated with DMSO or 500nM ACBI1 for 24h prior to expression of an empty vector (EV) or SS18::SSX1 transgene for additional 72h and co-treatment with DMSO or ACBI1 or Log2-Fold changes of RNA-seq of SS18::SSX relative to EV in DMSO or ACBI1 conditions at SS18::SSX1 target loci in KHOS-240S cells displaying up-regulation upon SS18::SSX transgene expression such as *IGF2*, *TPPP*, *MAFA*, *SHH*, *HES4* and *ONECUT3*.

ACBI1 depleted SMARCA4 within one day, while SS18::SSX was expressed robustly (**Figure 3B**). RNA-sequencing revealed that ACBI1 affected gene expression in KHOS-240S cells; however, a distinct cluster of genes upregulated by SS18::SSX was similarly induced in both DMSO and ACBI1 treated cells, indicating that at least some oncofusion-driven transcription can be initiated in the absence of BAF ATPase activity (**Figure 3C**). By examining known SS18::SSX target genes, we confirmed that fusion overexpression in both DMSO and ACBI1-treated cells induces their upregulation, whereas ACBI1-treated cells lacking the fusion fail to induce these genes (**Figure 3D, 3E**). To examine the chromatin landscape in parental KHOS-240S cells and assess how BAF depletion and SS18::SSX overexpression affect chromatin accessibility, we performed ATAC-sequencing. ATAC-seq at SS18::SSX target loci, previously identified by CUT&RUN^16^, revealed baseline chromatin accessibility in parental KHOS-240S cells. ACBI1 treatment alone reduced accessibility of target genes, however, SS18::SSX expression induced chromatin re-opening in BAF-depleted cells and further enhanced chromatin opening relative to BAF-intact cells. Together, these findings demonstrate that the fusion can drive chromatin remodeling and transcriptional activation independently of BAF catalytic function (**Figure 3F, 3G**). This is consistent with our previous data showing that rare variant synovial sarcoma fusions, such as EWSR1::SSX and MN1::SSX, drive SSX-dependent transcription through alternative cofactor constellations that are not strictly or directly dependent on BAF complex activity^16,30^.

### Smarca4 deletion does not impair SS18::SSX-driven tumorigenesis in vivo

Mice that express SS18::SSX in mesenchymal compartments develop tumors that faithfully recapitulate human synovial sarcoma at the histological and molecular levels^9,16,30-33^. Since our data suggest that the BAF catalytic activity is not essential for the transcriptional activation of Polycomb targets by the oncofusion, we next asked whether *Smarca4* deletion affects synovial sarcoma formation in mice. *Smarca4*-floxed mice were bred to mice carrying the conditional expression alleles for human *SS18::SSX1* (hSS1) or *SS18::SSX2* (hSS2), each from the *Rosa26* locus. Littermate cohorts of *Smarca4^wt/wt^*, *Smarca4^wt/fl^* and *Smarca4^fl/fl^* pups were injected with TATCre at 8 days of life to simultaneously induce fusion expression and Cre-loxP-mediated deletion of the ATPase-coding exons of *Smarca4* (**Figure 4A, 4B**). *Smarca4^fl/fl^* animals developed tumors driven by either SS18::SSX1 or SS18::SSX2, with significantly accelerated growth compared to *Smarca4^wt/wt^* mice (**Figure 4C, 4D**). Histological analysis confirmed that tumors across all genotypes exhibited canonical synovial sarcoma morphologies, including monophasic, biphasic, and poorly differentiated subtypes, with no loss of defining features upon *Smarca4* disruption (**Figure 4E**). While, residual ATPase activity might still be present from *Smarca2* compensation, deletion of *Smarca4* did not impair synovial sarcoma formation and instead significantly accelerated tumor growth in both the hSS1 and hSS2 models. This finding suggests that, contrary to prevailing models that emphasize an essential oncogenic role for SMARCA4-containing BAF complexes in synovial sarcoma, SMARCA4 may exert context-dependent tumor-suppressive functions in this disease potentially by regulating broader gene expression programs beyond those directly controlled by the fusion oncoprotein.

**Figure 4:**
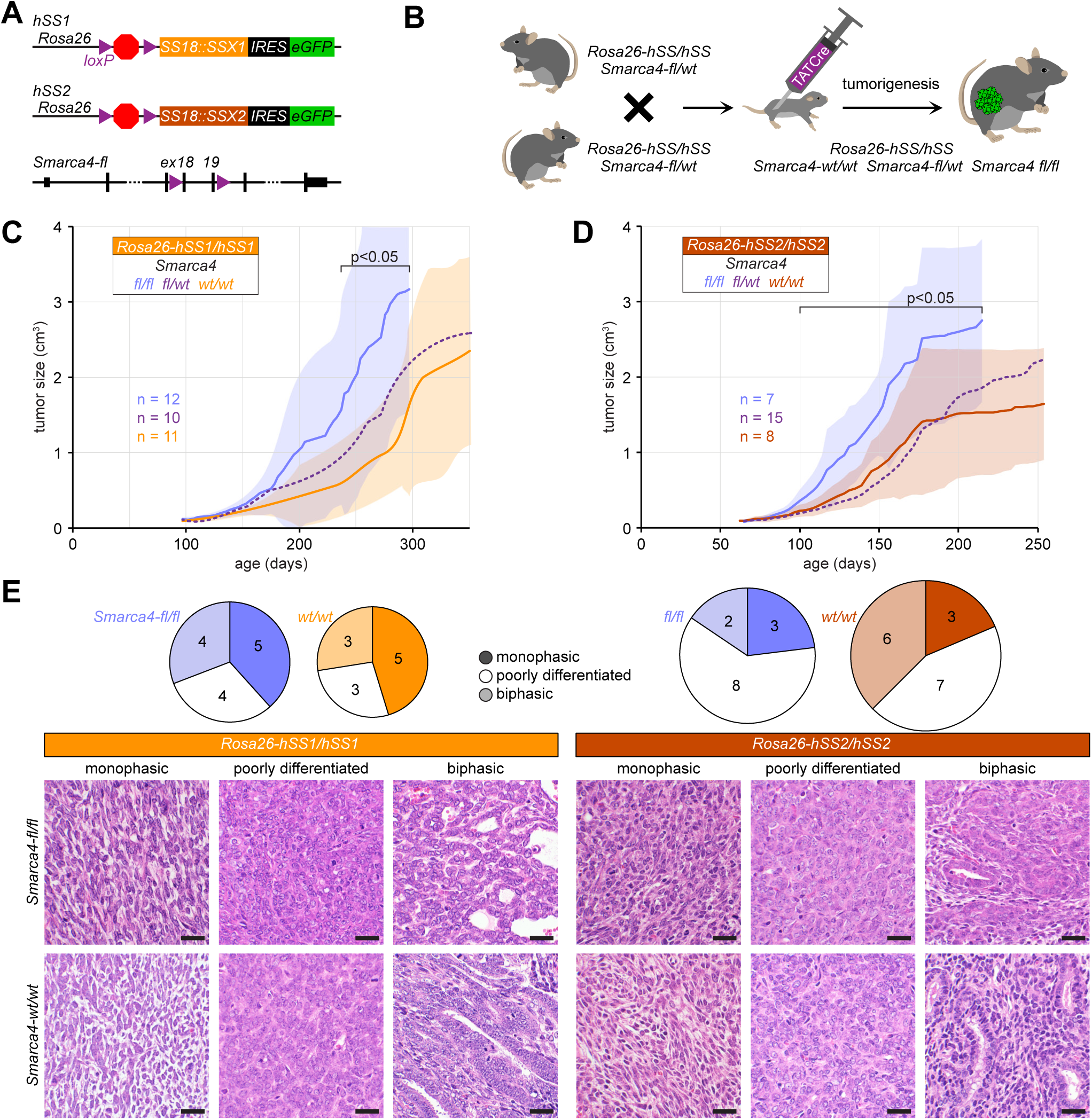
SS18::SSX-driven sarcomagenesis is preserved in *Smarca4*-deficient mice. **(A)** Schematic of hSS1 and hSS2 alleles that conditionally express SS18::SSX1 and SS18::SSX2 respectively from the Rosa26 locus upon Cre- mediated recombination of the loxP sites flanking a stop cassette and Smarca4-floxed allele that disrupts Smarca4 expression upon Cre- mediated recombination of the loxP sites flanking exons 18 and 19 of that gene, which code for critical residues in the ATPase site. **(B)** Schematic of the breeding strategy of heterozygous parents for Smarca4-fl, yielding littermate-controlled cohorts of homozygous wildtype and homozygous floxed pups that are injected in the hindlimb with TATCre protein at age 8 days to induce tumorigenesis that is followed with calipers. **(C)** Growth curves of mean caliper determined tumor size with shadows representing ± standard deviation for hSS1 tumors with variable Smarca4 genotypes. **(D)** Similar growth curves for hSS2 tumors. **(E)** The distribution of the three canonical histomorphology subtypes among the four listed genetic groups of tumors, as well as photomicrographs of hematoxylin and eosin stained histological sections of each.

### SS18 C-terminus recruits co-activators and induce transcription independently of BAF

SS18 contains an evolutionarily conserved domain near its N-terminus (SNH) that mediates stable incorporation into BAF complexes through direct interactions with core complex members, including the ATPase subunit SMARCA4^34-36^. The C-terminal region of SS18 is intrinsically disordered and enriched in glutamine, proline, glycine, and tyrosine residues (QPGY domain), characteristic of low-complexity transcriptional activation domains^37,38^. This QPGY-rich domain has been shown to activate transcription when fused to heterologous DNA-binding domains and may facilitate interactions with coactivators and other regulatory factors (**Figure 5A**). While direct interaction with BAF has been mapped to the SNH region, the intrinsically disordered QPGY domain has been shown to have transcriptional activation and phase separation properties^34-38^. To dissect how these domains, contribute to SS18::SSX transcriptional function, we expressed GFP-tagged SS18::SSX1 full-length (FL), an SNH-deleted mutant (ΔSNH), and a QPGY-deleted mutant (ΔQPGY), each fused to eGFP, from a lentiviral pLV backbone in KHOS-240S cells, to achieve strong and comparable expression across constructs (**Figure 5B**). Immunoprecipitation of eGFP followed by mass spectrometry showed that the ΔSNH mutant loses interaction with nearly all BAF subunits. Deletion of QPGY also reduced binding of several BAF subunits, but the effect was less pronounced than for ΔSNH (**Figure 5C-F**). We next asked to what extent the two mutants retain the ability to induce gene activation. RNA-seq in KHOS-240S cells showed that ΔSNH reduced the magnitude of fusion-induced transcriptional activation when compared with FL control, but still induced expression of the majority of SS18::SSX target genes compared with the eGFP control. In contrast, deletion of the QPGY domain nearly abolished transcriptional induction of these genes, consistent with the established role of the QPGY region as a potent transactivation module (**Figure 5G, 5H**). Notably, the median log₂ fold change of differentially expressed genes in ΔSNH-expressing cells was approximately 1.5, indicating substantial transcriptional induction despite strong diminution of the BAF interaction (**Figure 5I**). Comparative analysis of the ΔSNH and ΔQPGY interactomes identified transcriptional activators, including RBM14 (CoAA) and TAF2, as strongly enriched in the ΔSNH interactome relative to that of ΔQPGY (**Figure 5J**), consistent with a transactivation role for ΔSNH even following decreased association with the BAF complex. The nuclear co-activator RBM14 in particular, has been previously shown to interact with both SS18 and EP300, and can contribute to phase-separated coactivator assemblies at active regulatory elements^39-42^. Consistently with these observations, we detected RBM14 in SS18::SSX IP-MS datasets upon BAF depletion, and AlphaFold3 interaction predictions yielded ranking scores in the range of other known interactors (**Figure S3A-E**). Moreover, recent studies demonstrated that the QPGY domain of the fusion promotes phase separation, resulting in enrichment of EP300 at condensates^43^ and co-recruitment of SMARCA4 to these same structures^37,44^. EP300 detection in prior SS18::SSX pulldown experiments across multiple cellular backgrounds and conditions proved challenging^12^. This may relate to phase-separation-mediated interactions that are disrupted during cell lysis. To circumvent potential post-lysis interactome changes, we exploited a cellular strategy based on proximity labeling via the efficient biotin ligase miniTurbo (mT). This approach enables the capture of dynamic changes in proximity of a mT-tagged bait protein by streptavidin purification of covalently biotinylated proteins^45,46^. Transient expression of mT-SS18::SSX2 in HEK293T cells allowed us to map the proximal interactome of the fusion protein, revealing strong functional enrichment for transcriptional regulation and chromatin organization (**Figure S3F**). As expected, multiple BAF complex members were detected; notably, EP300 emerged as one of the top biotinylated proteins, and RBM14 was also significantly enriched (**Figure 5K**).

**Figure 5:**
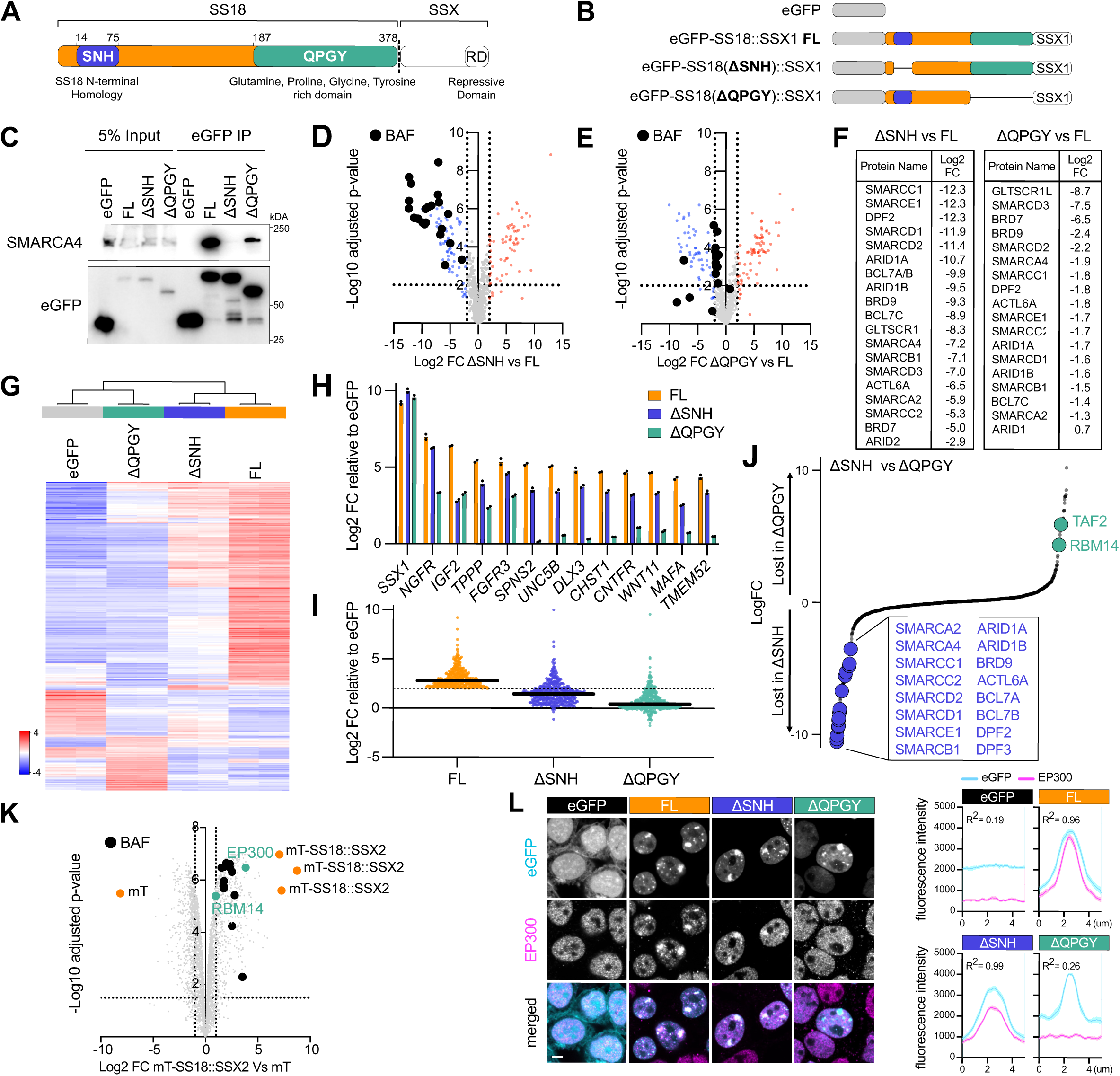
SS18 C-terminus engages co-activators and activates transcription without BAF. **(A)** Schematic of SS18::SSX displaying SS18 (orange) main domains, SS18 N-terminal homology (SNH, purple) and Glutamine, Proline, Gylcine, Tyrosine rich domain (QPGY, green). **(B)** Schematic of eGFP-fused constructs, eGFP alone, wild-type SS18::SSX isoform 2 (FL), SS18::SSX lacking SNH domain (ΔSNH) or SS18::SSX lacking QPGY domain (ΔQPGY). **(C)** Co-immunoprecipitation (co-IP) pulling down on eGFP in KHOS-240S cells expressing eGFP constructs representing one replicate. **(D), (E)** Volcano plots of mass spectrometry data following eGFP pull down of ΔSNH (E) or ΔQPGY (F) enrichment normalised to SS18 wild-type (FL). Data represents log2 fold change enrichment plotted on the x-axis and -log10 (adjusted p value) plotted on the y-axis. Blue/Red dots indicate lost/gained interactors relative to SS18::SSX1 isoform 2 (FL). Black dots indicate BAF members. n = 4 biological replicates. **(F)** List of BAF members displaying decreased interaction and their associated log2 fold change value in ΔSNH (left) or ΔQPGY (right) pull downs normalised to wild-type SS18::SSX1 isoform 2 (FL). Listed members are shown as black dots in (D) and (E). **(G)** RNA-seq heatmap showing the 1,000 most variable genes with a cutoff z-score of 4 from KHOS-240S cells expressing eGFP, FL, ΔSNH or ΔQPGY. n = 2 biological replicates. **(H)** Log2-transformed fold change of FPKM values in KHOS-240S RNA-seq relative to eGFP over selected SS18::SSX target genes. Data represents the mean. **(I)** Scatter plot of log2-transformed fold change of FPKM values in KHOS-240S RNA-seq relative to eGFP over genes up-regulated by SS18::SSX1 expression and showing >2 fold up-regulation. Biological replicates are combined using their mean. Bold line represents median and dotted line represents 2 cutoff. **(J)** Log2 fold change plot between ΔSNH and ΔQPGY mass spectrometry data following eGFP pull down in KHOS-240S cells. **(K)** Volcano plot of miniTurbo-v5-SS18::SSX2 proximity proteomics in HEK293T cells (miniTurbo = mT). Data represents log2 fold change enrichment plotted on the x-axis (mT-v5-SS18::SSX2 / mT-control)and -log10 (adjusted p value) plotted on the y-axis. Black dots indicate BAF members. EP300 and RBM14 are highlighted in green; mT and mT-SS18::SSX2 are highlighted in orange. n = 3 biological replicates. **(L)** Left: Immunofluorescence of HEK293T cells expressing eGFP, FL, ΔSNH or ΔQPGY (cyan) stained for EP300 (magenta). Images are representative of n = 2 biological replicates. Scale bar, 5 μm. Right: Profile of fluorescence intensity of eGFP, FL, ΔSNH or ΔQPGY over eGFP foci. Data represent the mean ± S.E.M of n = 2 biological replicates. For each replicate and condition, > 10 foci where profiled. For eGFP quantification in (N), random profiles across nucleus where generated. R² indicates Pearson correlation coefficient.

We have previously shown that SS18::SSX binds H2Aub-enriched nuclear foci in HEK293T cells via the SSX C-terminus, providing a tractable system to monitor recruitment of additional proteins to these foci^16^. First, we examined whether ΔSNH, and ΔQPGY mutants are also recruited to H2Aub foci in HEK293T cells. All three constructs were robustly targeted to H2Aub foci, indicating that, like full length SS18::SSX, both mutants retain the ability to localize at H2Aub-enriched regions (**Figure S3G**). Consistent with the proximity labeling identification of an association with EP300 (Figure 5K), we observed co-recruitment of EP300 at SS18::SSX/H2Aub foci. Importantly, the recruitment was preserved in the ΔSNH mutant, but lost upon deletion of the QPGY domain (**Figure 5L**). Together, our observations support a model in which the SS18 QPGY domain engages EP300-containing co-activator assemblies, even when direct BAF binding via SNH is disrupted.

### SS18::SSX relies on EP300/CREBBP to drive aberrant gene expression

Since EP300 is an important transcriptional activator that can be recruited by the ΔSNH mutant of SS18::SSX with diminished direct BAF association, we sought to explore its role in synovial sarcoma transcriptional activation. We previously showed that SS18::SSX chromatin binding is independent of the BAF ATPase activity^16^. HEK293T cells expressing FL SS18::SSX1 were treated with the dual EP300/CREBBP degrader dCBP-1^43^, resulting in efficient loss of EP300 but no disruption of SS18::SSX localization to H2Aub foci, indicating that neither EP300 nor CREBBP is required for fusion chromatin targeting (**Figure S4A**). Consistent with this, the core BAF subunit SMARCC1 also remained following dCBP-1 treatment. In contrast, ACBI1 treatment led to a robust depletion of SMARCC1 confirming disruption of BAF integrity (**Figure S4B**). Importantly, EP300 signal at SS18::SSX foci persisted despite ACBI1 treatment, indicating that oncofusion-mediated EP300 recruitment to repressed H2Aub foci can occur independently of BAF ATPase activity (**Figure 6A**).

**Figure 6:**
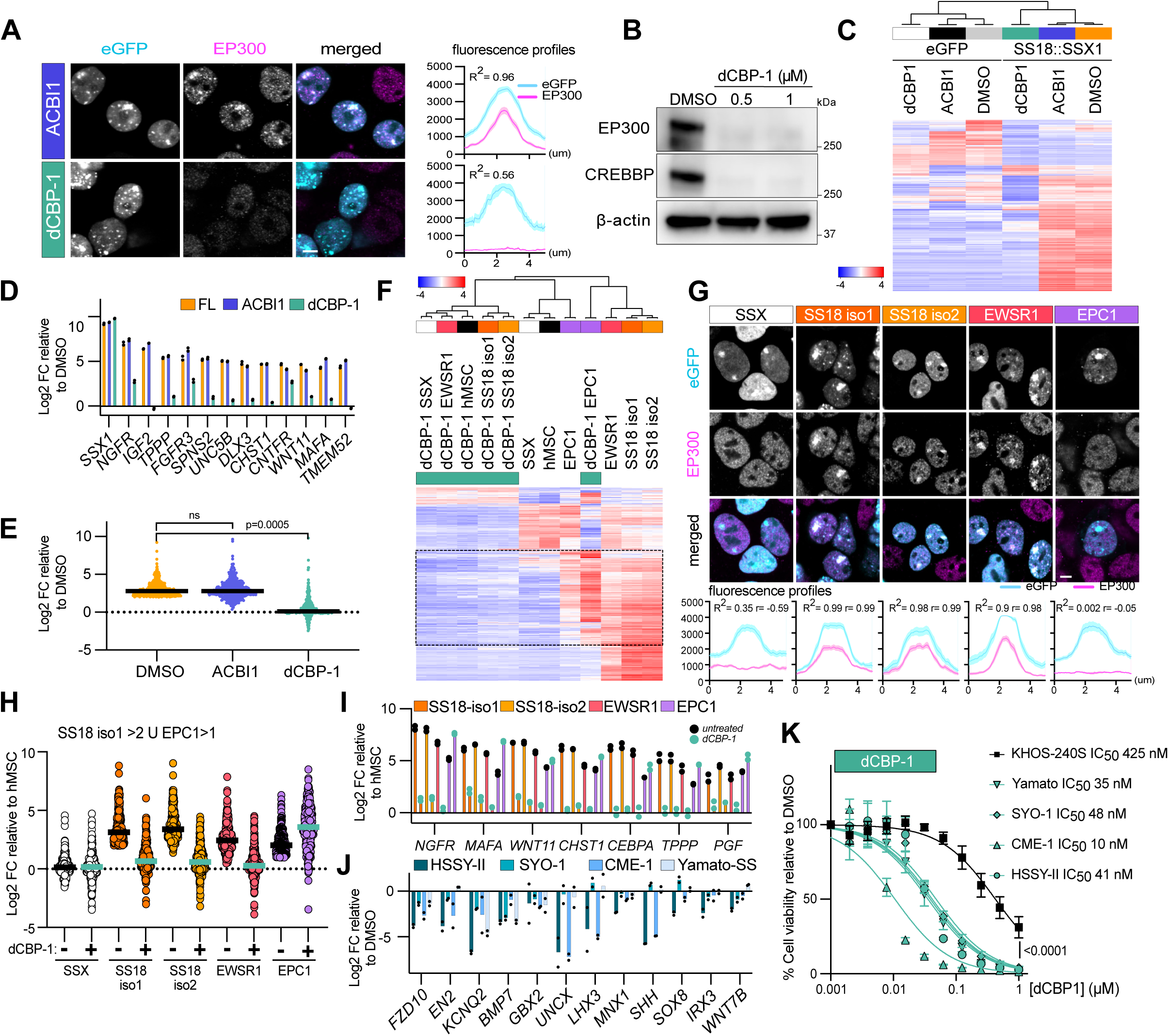
EP300/CREBBP engagement by SS18::SSX dictate its aberrant transcriptional program. **(A)** Left: Immunofluorescence of HEK293T cells expressing eGFP or wild-type SS18::SSX1, either untreated (FL) (cyan) or treated for 72h with 500nM ACBI1 or 500nM dCBP-1 stained for EP300 (magenta). Images are representative of n = 2 biological replicates. Scale bar, 5 μm. Right: Profile of fluorescence intensity. Data represent the mean ± S.E.M of n = 2 biological replicates. For each replicate, > 10 foci where profiled. R² indicates Pearson correlation coefficient. **(B**) Western Blot of KHOS-240S whole cell extracts treated for 24h with DMSO, 500nM or 1uM dCBP- 1 revealed using EP300, CREBBP or β-actin antibodies. **(C)** RNA-seq heatmap showing the 1,000 most variable genes with a cutoff z-score of 4 from KHOS-240S cells expressing eGFP or SS18::SSX1 (FL) treated for 72h with either DMSO, 500nM ACBI1 or 500nM dCBP-1. n = 2 biological replicates. (**D)** Log2-transformed fold change of FPKM values in KHOS-240S RNA-seq relative to eGFP over selected SS18::SSX target genes. Data represents the mean. **(E)** Scatter plot of log2-transformed fold change of FPKM values in KHOS-240S RNA-seq relative to eGFP over genes up-regulated by SS18::SSX1 expression and showing >2 fold up-regulation. Biological replicates are combined using their mean. Bold line represents median and dotted line represents cutoff of 2. **(F)** RNA-seq heatmap showing the 1,000 most variable genes with a cutoff z-score of 4 from wild-type hMSC or cells expressing eGFP-SSX1, eGFP-SS18::SSX1 isoform1, eGFP-SS18::SSX1 isoform2, eGFP-EWSR1::SSX1 or eGFP- EPC1::SSX1 treated for 48h with 500nM dCBP-1. n = 2 biological replicates.**(G)** Top: Immunofluorescence of HEK293T cells expressing wild-type eGFP-SSX1, eGFP-SS18::SSX1 isoform1, eGFP-SS18::SSX1 isoform2, eGFP-EWSR1::SSX1 or eGFP-EPC1::SSX1(cyan) stained for EP300 (magenta). Images are representative of n = 2 biological replicates. Scale bar, 5 μm. Bottom: profile of fluorescence intensity of eGFP constructs over SSX-rich eGFP foci. Data represent the mean ± S.E.M of n = 2 biological replicates. For each replicate and condition, > 10 foci where profiled. R² and r indicate Pearson correlation coefficients. **(H)** Scatter plot of log2-transformed fold change of CPM values in hMSC RNA-seq relative to untreated hMSC over the union genes up-regulated by SS18::SSX1 isoform1 (log2FC >2) and up-regulated by EPC1::SSX1 (log2FC >1). Biological replicates are combined using their mean. Bold line represents median. **(I)** Log2-transformed fold change of CPM values in hMSC RNA-seq relative to untreated cells over selected SSX target genes. Data represents the mean. **(J)** qRT-PCR displaying log2- transformed fold change of mRNA levels normalised by GAPDH in HSSY-II, SYO-1, CME-1 and Yamato-SS cells treated for 72h with 500nM dCBP-1 relative to DMSO. Data represents the mean of n = 2 biological replicates. **(K)** Viability of 4 synovial sarcoma cell lines (Yamato-SS, SYO-1, CME-1, HSSY-II) and KHOS-240S osteosarcoma cells treated with various concentrations of dCBP-1. The symbols represent the mean ± S.E.M of n = 3 biological replicates, the solid line represent the nonlinear curve fitting and its associated IC50. p-value represents extra sum-of- squares F test between KHOS-240S and SYO-1 nonlinear regression curves.

To evaluate the effect of dCBP-1 treatment in induction of SS18::SSX responsive genes we performed RNA-seq in KHOS-240S cells expressing SS18::SSX1 (**Figure 6B**). Whereas, as expected, ACBI1 had minimal impact on fusion target gene expression, dCBP-1 abolished expression of SS18::SSX-induced genes (**Figure 6C-E**). dCBP-1 treatment of mesenchymal stem cells expressing SS18::SSX (isoform 1 and 2) or EWSR1::SSX1 led to similar effects, showing a profound inhibition of fusion-induced genes in the absence of EP300/CREBBP (**Figure 6F**). Interestingly, fusion of SSX1 to EPC1, a member of NuA4/TIP60 complex, that has been shown to activate Polycomb target loci in endometrial sarcomas harbouring EPC1::PH1 fusions^47^, also activated a set of genes common to SS18::SSX and EWSR1::SSX fusions. However, EPC1::SSX1 failed to recruit EP300 to H2Aub foci in HEK293T (**Figure 6G**). Accordingly, and unlike SS18::SSX or EWSR1::SSX, EPC1::SSX-induced genes were not inhibited by dCBP-1 treatment (**Figure 6F, 6H-I**). Together, these results suggest that although EP300/CREBBP is required for gene activation in the context of SS18::SSX and other fusion oncoproteins such as EWSR1::FLI1^48,49^, this dependency is not a general feature of aberrant transcription driven by chimeric transcription factors. Instead, reliance on EP300/CREBBP appears to be restricted to specific transcriptional activators, which has important implications for the future use of EP300/CREBBP inhibitors or degraders across sarcoma subtypes.

Lastly, we treated a panel of synovial sarcoma cell lines with dCBP-1 to evaluate its effect on expression of known synovial sarcoma signature genes and cell viability. Treatment for 72 h led to a pronounced downregulation of known fusion target genes (**Figure 6J**). Consistently, synovial sarcoma lines exhibited high sensitivity to dCBP-1, with IC50s ranging from 10-48 nM, when compared to KHOS-240S osteosarcoma cells (**Figure 6K**). Together, these data demonstrate that while BAF catalytic activity is not critical for SS18::SSX-activated transcription, EP300/CREBBP function is selectively required for SS18::SSX-driven gene activation and synovial sarcoma cell survival.

## DISCUSSION

Synovial sarcoma is widely considered to arise from aberrant BAF complex function. Incorporation of the SS18::SSX fusion into BAF complexes destabilizes canonical BAF (cBAF) integrity and enriches ncBAF assemblies at Polycomb-marked chromatin due to the affinity of the SSX C-terminal domain for the nucleosome acidic patch with H2AK119ub1^17-19^. Together, these compositional and targeting alterations create a complex regulatory landscape in which BAF activity is perturbed both through disruption of many of its normal functions at canonical enhancers and aberrant chromatin localization, resulting in activation of normally silenced developmental gene programs. This duality has complicated efforts to distinguish whether synovial sarcoma pathogenesis primarily reflects loss-of-function effects on BAF activity, gain-of-function properties conferred by the fusion, or a combination of both.

An assumption emerging from this model is that aberrant localization of BAF to Polycomb target genes is required for their inappropriate activation. Although BAF complexes are well established to play a conserved role in antagonizing Polycomb-mediated repression^50^ the extent to which BAF activity is required for SS18::SSX-driven transcriptional activation in synovial sarcoma has not been directly tested before. Here, we challenge this view by demonstrating that acute degradation of the BAF ATPase subunits SMARCA2 and SMARCA4, or loss of core BAF components, fails to deactivate SS18::SSX-dependent transcriptional programs or substantially impair synovial sarcoma cell viability. Instead, SS18::SSX retains the capacity to induce target gene expression at fusion-occupied loci, independently of BAF catalytic activity. Beyond the limited effect on direct SS18::SSX target genes, disruption of SMARCA2/4 could potentially amplify aspects of fusion-driven transcription by further destabilizing cBAF, a mechanism also implicated in synovial sarcomagenesis^18^. This is supported by our *in vivo* findings showing that SS18::SSX-driven tumorigenesis proceeds, and is even accelerated, in a *Smarca4*-deficient background. Importantly, and perhaps most surprising, the ability of SS18::SSX to activate its direct target genes is maintained in the absence of BAF interaction, as domain-mapping experiments demonstrate that deletion of the SNH domain, which mediates direct interaction with the BAF ATPase subunit^34-36^, only partially attenuates activation of many SS18::SSX target genes. In contrast, transcriptional activation of SS18::SSX-responsive genes is critically dependent on the C-terminal QPGY-rich domain of SS18.

Although BAF complexes facilitate chromatin accessibility, previous work has shown that rapid loss of BAF subunits results in both gene activation and repression, indicating that transcriptional induction is not uniformly dependent on BAF-mediated remodeling^2,51^. Emerging evidence also suggests context-dependent BAF function and compensatory mechanisms. While enhancers often remain durably repressed upon BAF loss, promoters frequently recover accessibility through alternative remodelers such as EP400/TIP60, restoring nucleosome positioning and transcription^52^. Our findings showing the ability of SS18::SSX to activate transcription in the absence of BAF are also supported by uterine tumors resembling ovarian sex-cord tumors (UTROSCT). These tumors harbor GREB1-SS18 fusions, which lack the N-terminal SNH domain required for BAF incorporation, yet retain oncogenic properties^53,54^. Together, this indicates that minimal SS18 domains retain intrinsic transcriptional activation capacity independent of BAF recruitment.

Several studies have demonstrated that targeted recruitment of EP300/CREBBP catalytic activity is sufficient to induce transcription at endogenous promoters and enhancers, including through deposition of H3K27ac, preinitiation complex assembly, and RNA polymerase II recruitment^55-57^. Additionally it was recently demonstrated that CREBBP can drive zygotic genome activation via catalytic-independent mechanisms that recruit and stabilize RNA Pol II^58^. Our data support a model in which SS18::SSX engages coactivators such as EP300 to sustain transcription independently of BAF remodeling enzymatic activity. Whether EP300 recruitment occurs primarily through direct interactions with the QPGY domain, via cooperation with additional transcription factors, or through condensate-based assembly remains an open question. Likewise, the extent to which other transcriptional regulators contribute to the maintenance of SS18::SSX-driven transcriptional activation in the absence of BAF activity remains to be determined. Notably, colocalization of H2Aub-bound SS18::SSX with activating chromatin features such as broad H3K4me3 has led to the proposal that MLL complex is required for expression of fusion targets which can be inhibited by WDR5 depletion^59,60^. Finally, it should be noted that the ability to activate or sustain transcription independently of BAF is likely to be gene- and context-dependent, and influenced by the epigenetic state of individual loci in the cancer cell of origin. Still, our results show that, in synovial sarcoma cells, the majority of SS18::SSX-activated genes do not require continuous BAF activity to sustain their expression and, in certain contexts, can be induced de novo in the absence of functional BAF remodeling.

Together, these findings support a revised model in which BAF retargeting alone is insufficient to explain fusion-driven gene activation and instead highlight coactivator-dependent transcription as a dominant and therapeutically exploitable mechanism in synovial sarcoma. In this updated model, SS18::SSX is repositioned alongside other sarcoma-associated fusion oncoproteins, including EWSR1::FLI1 (Ewing sarcoma)^48,49^, PAX3::FOXO1 (alveolar rhabdomyosarcoma)^61^, HEY1::NCOA2 (mesenchymal chondrosarcoma)^62^, and CIC::DUX4^63^, which all converge on EP300/CREBBP as a central coactivator vulnerability. Indeed, each of these fusions engages EP300/CREBBP through distinct activation domains to drive lineage-inappropriate transcriptional programs, and pharmacologic inhibition or degradation of EP300/CREBBP has been shown to suppress these programs across multiple sarcoma subtypes^48,49,61-63^. Future studies should delineate the contributions of additional coactivators or phase-separation mechanisms and explore combinatorial therapeutic strategies incorporating EP300/CREBBP degraders. Collectively, these observations position EP300/CREBBP-targeted therapies as a potential unifying treatment approach for fusion-driven sarcomas that depend on aberrant transcriptional activation.

## METHODS

### Cell culture

Human synovial sarcoma cell lines HS-SY-II (RRID: CVCL_8719), SYO-1 (RRID: CVCL_7146), Yamato-SS (RRID: CVCL_6C44) were obtained from their original source laboratories, while CME-1 was obtained from Stefan Fröhling’s laboratory in DKFZ. Human osteosarcoma KHOS-240S (RRID: CVCL_2544), human embryonic kidney HEK293T (RRID: CVCL_0063) and the human Mesenchymal Stem Cell-hTERT immortalized (hMSC) (ASC52telo, RRID:CVCL_U602) cell lines were purchased from the American Type Culture Collection (ATCC). HEK293GP cells used for retrovirus production were obtained from Takara Bio (631458). HS-SY-II, SYO-1, Yamato-SS, KHOS-240S, HEK293T and HEK293GP were cultured in DMEM (Gibco) while CME-1 was cultured in RPMI 1640 (Gibco), all supplemented with 10% fetal bovine serum (FBS) and penicillin-streptomycin. The hMSCs were cultured in MesenPRO RS Medium (Gibco, 12746-012) supplemented with L-glutamine (Sigma-Aldrich, G7513-100ML) at a final concentration of 2 mM. The SMARCA2/SMARCA4 degrader ACBI1 (2375564-55-7) and the EP300/CREBBP degrader dCBP-1 (2484739-25-3) were purchased from MedChemExpress, resuspended in dimethyl sulfoxide (DMSO) and kept at −80°C. Cells were treated for 24 h, 72 h or 96 h with 500 nM ACBI1 or 500 nM dCBP-1.

### Plasmids

The dual-sgRNA-eGFP-P2A-Puro-hU6-mU6 plasmid was cloned using as donor vector the original dual-sgRNA-CD90-hU6-mU6 (Addgene, 154194) digested with XhoI (NEB) and SalI-HF (NEB) and as insert the GFP-P2A-Puro-WPRE sequence PCR amplified from the pLKO.1-sgRNA vector. The assembly was designed and performed in a single step using NEBuilder HiFi DNA Assembly.

sgRNAs for CRISPR knockout were designed using the tool from Sanjana Lab (currently unavailable) and cloned as previously described^64^. In brief, sgRNAs were cloned by annealing and phosphorylating two DNA oligo pairs for site 1 (human U6-hU6) and site 2 (mouse U6-mU6) and ligating into an Esp3I (NEB) and BbsI-HF (NEB) digested dual-sgRNA-eGFP-P2A-Puro-hU6-mU6 plasmid. Transformation was carried into Stbl3 bacteria.

MSCV-PGK-Puro and MSCV-PGK-SS18::SSX1-Puro plasmids were taken from Banito, et.al, 2018^13^.

pLV-EF1α-eGFP-SS18-IRES-Neo, pLV-EF1α-eGFP-SS18::SSX1-IRES-Neo and pLV-EF1α-eGFP-SSX1-IRES-Neo were taken from Benabdallah, et.al 2023^16^, previously cloned into pLV-EF1α-IRES-Neo lentiviral backbone (Addgene, 85139) containing a neomycin selection cassette.

pLV-EF1α-eGFP-SS18(ΔSNH)::SSX1-IRES-Neo and pLV-EF1α-eGFP-SS18(ΔQPGY)::SSX1-IRES-Neo were generated from pLV-EF1α-eGFP-SS18::SSX1-FL-eGFP-IRES-Neo using Q5 site directed mutagenesis kit, utilizing NEB’s KLD enzyme mix reaction. Deletions of defined protein domains were generated by PCR using complementary oligonucleotides that flank the region to be deleted. PCR products were treated with a kinase-ligase-DpnI (KLD) mixture to phosphorylate and circularize the amplified product and to selectively degrade the methylated parental plasmid.

For cloning of the miniTurbo-V5-SS18::SSX2 plasmid, SS18::SSX2 was cloned by Gibson assembly (NEB, E5510S) into BamHI (NEB) digested pENTR221_miniTurbo_V5^46^. Primers to amplify SS18::SSX2 contained Gibson compatible overhangs. The pSB_EF1α_SS18::SSX2 vector was used as a template. The resulting plasmid pENTR221_miniTurbo_V5_SS18::SSX2 was subsequently cloned into a Gateway compatible destination vector using LR clonase (Thermo Fisher Scientific, 11-791-020) to generate pLenti_miniTurbo_V5_SS18::SSX2.

For the SSX1 fusion constructs, EWSR1, SS18 isoform1 and SS18 isoform2 were re-cloned from previous constructs^16,32^ into pLV-EF1α-eGFP-IRES-Hygro, originally obtained from Addgene (Addgene, 85134) and modified to contain the eGFP cassette and restriction enzyme sites between the eGFP and the IRES sequence to act as landing pads. EPC1 cDNA (NM_025209) was obtained from GenScript (EPC1_OHu20692D_pcDNA3.1+/C-(K)-DYK).

### Virus production and transduction

For lentivirus production, 1,5 x 10^6^ HEK293T cells were transfected with 3.5 μg of constructs and helper vectors (2.5 μg of psPAX2 and 0.9 μg of VSV-G). For retroviral infection, 2 x 10^6^ HEK293GP cells containing a gag-pol insertion were transduced with 3 μg of MSCV vectors and 0.9 μg of VSV-G. Transfection of packaging cells was performed using polyethyleneimine (Polysciences, 23966-2) by mixing with DNA in a 3:1 ratio. Medium was changed after 16 h and viral supernatants were collected 48 h after transfection, filtered through a 0.45-μm filter and supplemented with 4 μg/ml polybrene (Sigma) before adding to target cells. Downstream experiments using sgRNAs for knockouts were performed 7 days after sgRNA induction (Western Blots, TIDE assay, RNA-seq, ATAC-seq). Downstream experiments using overexpression of MSCV or pLV constructs (Western Blots, RNA-seq, ATAC-seq, Immunoprecipitation-Mass Spectrometry and immunofluorescence) were performed 48 or 72 h after induction (indicated in figure legends).

### Generation of Cas9 stable cell lines

HS-SY-II, SYO-1, Yamato-SS, CME-1 synovial sarcoma cell lines and KHOS-240S osteosarcoma cell line were transduced with lentiCas9-Blast^65^ (Addgene, 52962) and selected using 20 μg/ml blasticidin (Gibco) to generate stable Cas9-expressing cell lines. Cells were subsequently transduced with sgRNAs. After 2 days of infection, cells were selected with 2 µg/ml puromycin (Gibco).

### Whole cell protein extracts and western blotting

Cells grown in 6-well plates or 10 cm^2^ petri-dishes were collected and washed in PBS. Cell pellets were incubated with RIPA buffer (Cell Signalling Technology) supplemented with protease inhibitors (Roche) for 30 min and cleared by centrifugation (15 min, 16.000 rpm, 4°C). Protein lysates were quantified using a BCA Protein Assay (Pierce). Lysates were then denatured in 4X Laemmli (Bio-rad), 0.25% β-mercaptoethanol (Thermo Fischer) at 95°C for 5 min, then run in Mini-PROTEAN Precast Gels (Bio-Rad) and transferred onto membranes using Trans-Blot Turbo. PVDF membranes were blocked in 5% milk in TBST. Western blots were visualized using an Amersham Imager 680.

### Tracking of Indels by Decomposition (TIDE) Analysis

Genomic DNA was extracted from Yamato -SS Cas9 infected cells expressing sgRNA for 7 days and control cells infected with sgSAFE using DNeasy Blood & Tissue kit (QIAGEN) following manufacturer’s protocol. The region targeted with sgRNA was amplified and purified using PCR purification kit (QIAGEN). Following Sanger sequencing of the PCR amplicons, sequences were analyzed using the TIDE tool (http://shinyapps.datacurators.nl/tide/) to calculate the percentage of insertions and deletions and assess sgRNA efficiency.

### Cell Competition Assays

HS-SY-II, SYO-1, Yamato-SS, CME-1, and KHOS-240S cell lines with a constitutive expression of Cas9 were transduced with a sgSAFE or a plasmid containing sgRNA targeting the respective gene of interest. Infections were performed to obtain an infection efficiency of around 70-80%. sgRNA infected cells become GFP+ due to the eGFP reporter of the dual-sgRNA vector. The cells were then cultured over a period of 24 days, and the percentage of GFP+ cells was measured using a Fortessa FACS machine. Data was analyzed using Flowjo software (https://www.flowjo.com/).

### Drug treatments

Synovial sarcoma cell lines (HS-SY-II, SYO-1, CME-1, Yamato-SS) and control osteosarcoma cell line (KHOS-240S) were assessed for sensitivity to ACBI1 and dCBP-1. Cells were seeded into 384-well plates at the following densities: HS-SY-II (2,000 cells/well), SYO-1 (500 cells/well), and CME-1, Yamato-SS, and KHOS-240S (1,000 cells/well). Cells were treated with a 10-point dose-response curve (highest final concentration 1 μM, two-fold serial dilutions) and incubated for 5 days at 37°C and 5% CO₂. Cell viability was measured as ATP levels using CellTiter-Glo® Luminescent Cell Viability Assay (Promega) on a Tecan plate reader.

### RNA extraction and qPCR

RNA was prepared using the RNeasy Mini Kit (QIAGEN) according to the manufacturer’s protocol and including DNase I (QIAGEN) treatment. cDNA was synthesized from purified RNA with RevertAid Reverse Transcriptase (Thermo Scientific) primed with polyA tail and random hexamers. qPCR was carried on the Roche LightCycler 480 Real-Time PCR System using Power SYBR Green PCR Master Mix (Applied Biosystems). The real-time thermal cycler was programmed as follows: 15 min Hotstart; 44 PCR cycles (95°C for 15 s, 55°C for 30 s, 72°C for 30 s).

### RNA sequencing and analysis

HSSY-II cells were treated for 72 h with 500nM ACBI1 or DMSO or transduced with lentiviral constructs for *SS18::SSX1* knockout (sgSSX), combined knockout of *SMARCA2/SMARCA4* (sgSMARCA2/4), combined knockout of *SMARCC1/SMARCC2* (sgSMARCC1/2) or with sgSAFE as a control (Fig. 2A, B). KHOS-240S cells were treated for 24 h with DMSO or 500nM ACBI1 prior to expression of MSCV-SS18::SSX1 or MSCV-EV for additional 72 hs and in co-treatment with DMSO or ACBI1 (Fig. 3C, D, E). KHOS-240S cells were transduced with pLV-EF1α-eGFP, pLV-EF1α-eGFP-SS18::SSX1, pLV-EF1α-eGFP-SS18::SSX1(ΔSNH) or pLV- EF1α-eGFP-SS18::SSX1(ΔQPGY) for 72 h (Fig. 5G, H, I). KHOS-240S cells were treated for 24 h with DMSO, 500nM ACBI1, 500nM dCBP-1 prior to expression of pLV-EF1α-eGFP, pLV-EF1α-eGFP-SS18::SSX1 for additional 72 h and in co-treatment with DMSO, ACBI1 or dCBP-1 (Fig. 6C, D, E). Total RNA was isolated using Qiagen RNeasy kit (74004). Library preparation and sequencing was performed by the DKFZ Next Generation Sequencing (NGS) core facility on NovaSeq 6000 (paired-end 100 bp, S4 flow cell). The data was demultiplexed and aligned to the reference human genome (HG19) by the DKFZ NGS core facility. Data shows FPKM values.

RNA sequencing of SSX chimeric fusions in figure (6F, H, I) was done using Plasmidsaurus RNA sequencing service. Briefly, 72 h after transduction and 48h after 500nM dCBP-1 treatment, 200000 cells per replicate were lysed in 50ul of Zymo DNA/RNA shield (R1100-50) and sent for sequencing on Illumina platform using a 3’ end counting approach. Details can be found here (https://plasmidsaurus.com/technical-documentation/rna). Data shows CPM values of samples in HG38.

FPKM or CPM values were used as input for iDEP (http://bioinformatics.sdstate.edu/idep93/), an online tool which uses R-Packages for RNA analysis to generate heatmaps of the most variable genes and perform differential expression analysis.

### ATAC sequencing

HSSY-II cells were treated for 3 days with 500nM ACBI1 or vehicle and transduced for 7 days with sgSSX or sgSAFE. KHOS-240S cells were treated for 24 hs with DMSO or 500nM ACBI1 or and then transduced with MSCV-SS18::SSX1 or MSCV-EV while in co-treatment with DMSO or ACBI1 for 72 h. Cells were used for downstream ATAC sequencing analysis. ATAC sequencing libraries were prepared using the Active Motif Pre-Indexed Assembled Tn5 Transposomes (Cat. No. 53152) from 75,000 cultured cells per replicate, following the manufacturer’s protocol with minor modifications. Cells were pelleted at 500 × g for 5 min at 4°C, washed with ice-cold 1× PBS, and resuspended in ice-cold ATAC lysis buffer for 10 min on ice. Nuclei were pelleted at 500 x g for 10 min at 4°C. Nuclei were resuspended in 46 μL tagmentation mix (25 μL 2 x tagmentation buffer, 2 μL 10 x PBS, 0.5 μL 1% digitonin, 0.5 μL 10% Tween-20, 18 μL nuclease-free water) and added to wells containing 4 μL pre-indexed Tn5 transposomes (8 μM). Reactions were incubated at 37°C with shaking (800 rpm) for 50 min, stopped with 1.8 μL 0.5 M EDTA and 0.5 μL 10% SDS, and heated at 55°C for 3 min. Tagmented DNA was purified using 1.2 x AMPure XP beads, washed twice with 200 μL 80% ethanol, and eluted in 22 μL 0.1 x Tris EDTA (TE) buffer. Libraries were amplified in 50 μL reactions (20 μL DNA, 2.5 μL each 25 μM P5/P7 primers, 25 μL NEBNext Ultra II Q5 Master Mix) using: Incubation at 72°C 5 min and at 98°C 30 sec followed by 8 cycles of 98°C 10 s, 63°C 30 s, 72°C 1 min. Products were purified with 1.2 x AMPure XP beads and eluted in 18 μL 0.1 x TE buffer. Libraries were validated using the Agilent High Sensitivity kit and then normalized to 6 nM, pooled and size-selected using double-sided AMPure XP beads (0.6 x right-side, 0.9 x adjusted left-side selection). Pools were sequenced on NovaSeq 6000 (paired-end 100 bp, S4 flow cell).

### ATAC sequencing analysis

ATAC-seq reads were analyzed using the Galaxy platform of DKFZ. FASTQ files were quality-checked with FastQC and MultiQC, trimmed with TrimGalore! and mapped to CHM13_T2T_v2.0 using Bowtie. Duplicate reads were marked and removed, filtered for mapping quality ≥20, and mitochondrial reads were excluded. Filtered BAM files were converted to BED format using bedtools bam to bed. Peaks were called with MACS2 callpeak (no model, shift -100, extension size 200, q=0.05) to account for Tn5 transposase bias, generating narrowPeak and bedGraph files. BedGraph files were converted to bigWig for visualisation of gene tracks in the UCSC Genome Browser (https://genome.ucsc.edu). Log2FC files were generated using bigwigcompare. For heatmaps and metaplot profiles, read densities of the various conditions were centered around peak signals with a ±5-kilobase (kb) window from peak center and binned with 50 bp using the computeMatrix and plotHeatmap functions from deepTools.

### ChIP and CUT&RUN analysis

For HSSY-II ChIP data, SMARCA4 input (SRR6451585) and IP (SRR6451598) as well as SS18::SSX1 (HA) input (SRR6451607), SS18::SSX1 (HA) IP (SRR6451595) were obtained from deposited GEO under the accession number GSE108929. For KHOS-240S CUT&RUN data, SS18::SSX profile (SRR24728103) was obtained from GSE205955. GREAT tool was used to define associated genes of SMARCA4-enriched ChIP-seq signal (http://bejerano.stanford.edu/great/public/html/index.php) using default settings.

### Co-immunoprecipitation

For endogenous co-immunoprecipitation of SS18::SSX1 and immunoprecipitation submitted to mass spectrometry analysis, approximately 5 x 10^7^ HSSY-II or KHOS-240S cells treated for 72 h with DMSO or 500 nM ACBI1 were harvested and nuclear extracts were prepared using a Nuclear Complex Co-IP Kit (Active Motif, 54001) with high-stringency IP and wash buffers, lacking added NaCl or detergent, followed by enzymatic DNA shearing for 1.5 h at 4°C. Nuclear extracts were incubated overnight at 4°C with either anti-SS18::SSX1 antibody or isotype-matched IgG control (3 μg antibody per reaction), with 10% of the extract retained as input. The following day, Protein A magnetic beads (Dynabeads Protein A, Thermo Fisher) were added and incubated for 1.5 h at 4°C with rotation, washed on a magnetic rack in high-stringency IP buffer, and eluted in RIPA/Laemmli sample buffer containing β-mercaptoethanol at 95°C for 10 minutes.

For immunoprecipitation and immunoprecipitation submitted to mass spectrometry analysis, approximately 5 x 10^7^ KHOS-240S cells transduced with pLV-EF1α-eGFP-SS18::SSX1-FL-eGFP-IRES-Neo, pLV-EF1α-eGFP-SS18(ΔSNH)::SSX1-IRES-Neo, pLV-EF1α-eGFP-SS18(ΔQPGY)::SSX1-IRES-Neo or pLV-EF1α-eGFP-IRES-Neo were collected and washed twice in ice-cold PBS. Whole cell extracts were generated by resuspending the pellets in co-IP buffer (100 mM NaCl, 10% glycerol, 50 mM Tris-HCl pH 8, 1% Triton X-100, 2 mM MgCl2, 0.1 mM ZnCl2, 0.2 mM EDTA, 1 mM DTT, EDTA-free protease inhibitors) supplemented with Benzonase nuclease and incubated for 1 h at 4°C. Lysates were cleared by centrifugation (10 min, 14,000 g, 4°C) and incubated with GFP-Trap Magnetic Agarose beads (ChromoTek) for 1.5 h at 4°C. Beads were washed three times with co-IP buffer and eluted in Laemmli/RIPA sample buffer containing β-mercaptoethanol at 95°C for 10 minutes.

For immunoprecipitation and immunoprecipitation submitted to mass spectrometry analysis approximately 10^7^ HEK293T cells transfected with pLV-EF1α-eGFP-SS18-IRES-Neo, pLV-EF1α-eGFP-SS18::SSX1-IRES-Neo and pLV-EF1α-eGFP-SSX1-IRES-Neo were collected aÅer 4 days of overexpression and 3 days treatment with 500nM ACB1 or DMSO. Cells were lysed with 150mM NaCl, 50mM Tris (pH 7.5), 5% glycerol, 1% IGPAL-CA-630, protease inhibitors (EDTA-free), phosSTOP, 1mM MgCl2 and 1% Benzonase. Lysates were incubated on ice for 20 min and centrifuged at 14000 rpm at 4°C for 10 min. 1mg lysate was incubated with GFP-Trap Magnetic Agarose beads (ChromoTek) for 1.5 h at 4°C. Samples were washed three times with wash buffer (150mM NaCl, 50mM Tris (pH 7.5), 5% glycerol, 0.05% IGPAL-CA-630) and further washed three times with basic wash buffer (150mM NaCl, 50mM Tris (pH 7.5), 5% glycerol). The beads were washed with 50mM ammonium bicarbonate (ABC) buffer. 100µl 4M urea in 50mM ABC with 1mM dithiothreitol and 100µl 10mM chloroacetamid were added to the beads which were shaken for 20 min 700 rpm. The beads bound proteins were digested overnight at 37 °C with shaking (1700 rpm) using 1ug trypsin and LysC. Digestion was stopped by adding 2µl trifluoroacetic acid (TFA). Peptides were desalted on 2 layers of C18 StageTips and dried in a vacuum concentrator at 45 °C and reconstituted in buffer A* (2% acetonitrile, 0.1% TFA). Peptide concentration was determined using a Nanodrop spectrophotometer and samples were stored at -20°C until measurement. Eluates and input samples were analyzed by SDS-PAGE and immunoblotting as described above.

### Proximity labeling in intact cells

10cm^2^ culture plate of HEK293T cells (50% confluent) was transfected with 3.4μg of pLenti_miniTurbo_V5 or pLenti_miniTurbo_V5_SS18::SSX2 using PEI (Polysciences, 24765). 48h after transfection, 500μM biotin was added for 10min. Labeling was stopped by transferring the cells to ice and washing 3 times with ice-cold PBS. Cells were collected and lysed in 600μL of lysis buffer (50mM Tris-HCl pH 7.5, 125mM NaCl, 5% glycerol, 0.2% NP-40, 1.5mM MgCl_2_ and protease inhibitors). After 30min incubation at 4°C, lysates were clarified by centrifuging at 15,000g for 20min.

### Enrichment of biotinylated proteins and on-bead digestion

For streptavidin pull-down of the biotinylated proteins, 200μg of protein was incubated with 50μL of lysis buffer-washed streptavidin magnetic beads (Thermo Fisher Scientific, 11205D) overnight at 4°C on a rotator. Beads were pelleted using a magnetic rack and each bead sample was washed three times with lysis buffer and three times with 50 mM ammonium bicarbonate pH 8.0. Samples were stored in 100 μL 50 mM ammonium bicarbonate pH 8.0 at -80°C prior to analysis. Tryptic digestion was performed directly on beads by incubating them with 1 μg of trypsin in a total volume of 300 μL 50 mM NH4HCO3 (150 μL were added to the initial 150 μL volume) at 37°C overnight. Additional 0.5 μg of trypsin were added to the samples and incubated for 2h at 37°C. Beads were pelleted by centrifugation at 2,000 g for 5 min, and the supernatant was transferred to a fresh Eppendorf tube. Beads were washed once with 100 μL of 50 mM ABC buffer, and these washes were pooled with the first supernatant. Complete separation of the beads was performed with a magnetic separation rack. FA was added to the eluates to a 1% final concentration. Samples were cleaned up through C18 tips (polyLC C18 tips) and peptides were eluted with 80% ACN, 1% TFA in water. Next, samples were diluted to a final concentration of 5% ACN and 0.1% TFA and loaded into SCX columns (polyLC). Peptides were eluted in 5% NH4OH, 30% methanol. Finally, samples were evaporated to dryness in the Speed Vac, reconstituted in 50 μL 3% ACN, 1% FA aqueous solution, and diluted 1:4 in 0.1% FA aqueous solution for MS analysis.

### Mass Spectrometry-Analysis

Eluted proteins from the co-IP experiments of HSSY-II cells (Figure 1B, 1C) and KHOS-240S (Figure 5D-F) were submitted for mass spectrometry at the DKFZ Proteomics Core facility. 45 µl of the IP eluates (treated as 5 ug) were digested (trypsin) using an AssayMAP Bravo liquid handling system (Agilent technologies) running the autoSP3 protocol according to Müller et, al. 2020^66^. Eluting peptides were analyzed in the mass spectrometer using data dependent acquisition (DDA) mode. A full scan at 60k resolution (380-1400 m/z, 500% AGC target, 100 ms maxIT) was followed by up to 1.5 seconds of MS/MS scans. Peptide features were isolated with a window of 1.2 m/z, fragmented using 26% NCE. Fragment spectra were recorded at 15k resolution (100% AGC target, 150 ms maxIT). Unassigned and singly charged eluting features were excluded from fragmentation and dynamic exclusion was set to 10 s. Each sample was followed by a wash run (20 min) to minimize carry-over between samples. Instrument performance throughout the course of the measurement was monitored by regular (approx. one per 48 h) injections of a standard sample and an in-house shiny application.

Data analysis of the KHOS-240S cell mass spectrometry data (Figure 5D-F) was carried out by MaxQuant (v2.1.4.0)^67^ using an organism specific database extracted from Uniprot.org (human reference database with one protein sequence per gene, containing 20,650 unique entries from 21st of January 2025, EGFP and SS18::SSX1 sequence). Settings were left at default with the following adaptions. Match between runs (MBR) was enabled to transfer peptide identifications across RAW files based on accurate retention time and m/z. Fractions were set in a way that MBR was only performed within replicates. Label free quantification (LFQ) was enabled with default settings. The iBAQ-value^68^ generation was enabled. For the statistical analysis log2 transformed iBAQ values were used. The following steps were performed per statistical contrast and not across the whole data matrix. Protein groups with valid values in 70% of the samples of at least one condition were used for statistics. Variance stabilization normalization was applied to normalize across samples^69^. Adapted from the Perseus recommendations^70^, missing values being completely absent in one condition, were imputed with random values drawn from a downshifted (2.2 standard deviation) and narrowed (0.3 standard deviation) intensity distribution of the individual samples. For missing values with no complete absence in one condition the R package missForest was used for imputation^71^. The statistical analysis was performed with the R-package “limma”^72^, single contrasts were adapted from the user guide chapter *Two Group*. Alongside the first author of limma comments, the setup was adapted for proteomics data via setting the *eBayes* options *trend* and *robust* to *TRUE*. The p-values were adjusted with the Benjamini–Hochberg method for the multiple testing^73^.

Data analysis of the HSSY-II mass spectrometry data (Figure 1B, 1C) was carried out by MaxQuant^67^ using an organism specific database extracted from Uniprot.org (human reference database with one protein sequence per gene, containing 20,650 unique entries from 21^st^ of January 2025, SS18::SSX1 sequence). Settings were left at default with the following adaptions. Match between runs (MBR) was enabled to transfer peptide identifications across RAW files based on accurate retention time and m/z. Fractions were set in a way that MBR was only performed within replicates. Label free quantification (LFQ) was enabled, with the only within parameter groups option. The iBAQ-value^68^ generation was enabled. For the statistical analysis log2 transformed iBAQ values were used. The following steps were performed per statistical contrast and not across the whole data matrix. Protein groups with valid values in 70% of the samples of at least one condition were used for statistics. No normalization was applied. Adapted from the Perseus recommendations^70^ missing values being completely absent in one condition, were imputed with random values drawn from a downshifted (2.2 standard deviation) and narrowed (0.3 standard deviation) intensity distribution of the individual samples. For missing values with no complete absence in one condition the R package missForest was used for imputation^71^. The statistical analysis was performed with the R-package “limma”^72^, single contrasts were adapted from the user guide chapter Two Groups, multi conditions from Interaction Models. Alongside the first author of limma comments, the setup was adapted for proteomics data via setting the eBayes options trend and robust to TRUE. The p-values were adjusted with the Benjamini–Hochberg method for the multiple testing^73^.

For mass spectrometry analysis of HEK293T cells treated with 500nM ACBI1 or DMSO following eGFP-SS18::SSX1, eGFP-SS18 or eGFP-SSX pulldown (Figure S3A-D), 200ng peptides were loaded onto a nanoElute UHPLC system (Bruker Daltonics) coupled to a timsTOF Pro mass spectrometer via a CaptiveSpray nano-electrospray ion source. Separation was performed on a 25 cm IonOpticks Aurora column using buffer A (0.1% formic acid, 2% ACN) and buffer B (0.1% formic acid in 99.9% ACN). Peptides were separated with a 120-min gradient: 2–12% B (0–60 min), 12–20% B (60–90 min), 20–30% B (90–100 min), 30–85% B (100–110 min), followed by a 10-min wash at 85% B at a flow rate of 0.3 ul/min. TIMS elution voltages were calibrated using Agilent ESI-Low Tuning Mix (ions m/z 622, 922, 1222)). All measurements were conducted in DDA-PASEF mode with a recording window from m/z 100 to 1700 and a dimension range of 1/K0 0.65 to 1.40. Each top N acquisition cycle included 10 PASEF MS/MS scans. Precursor ions were selected for MS/MS based on an intensity threshold of 1000 a.u. and re-sequenced until reaching a threshold of 10,000 a.u. TIMS operated within a scan range of 100–1700 m/z, ramp time 100 ms, duty cycle 100%, cycle time 100 ms and spectra acquisition rate 9.43 Hz. Raw DDA MS data were processed using MaxQuant (v2.1.4.0)^67^ with the Andromeda search engine against the UniProt Human reference FASTA (downloaded on 7th March 2021). Default seàngs were applied, allowing a maximum of 2 missed cleavages. Proteins were assigned to the same protein group if two proteins could not be distinguished by unique peptides. Label-free quantification (LFQ) was performed using the MaxLFQ algorithm, with "match between runs" enabled to identify peptides across runs. iBAQ was switched on. The minimum ratio count for protein quantification was set to 2. Output data from Maxquant were imported into Perseus (v1.5.2.11)^70^. Intensity values were log2-transformed and missing values were imputed. Samples were normalized to SSX (DMSO and ACBI1). Statistical analyses were conducted using two-sample t-tests with a permutation-based FDR of 5%.

miniTurbo-related mass spectrometry (Figure 5K, S3F) was performed on an Orbitrap Eclipse Tribrid mass spectrometer (ThermoFisher Scientific, San Jose, CA) connected to an EVOSEP One (EVOSEP, Odense, Denmark) via nanoEasy Spray Source interface with a stainless-steel emitter (EV-1086 EVOSEP). Tryptic peptides were loaded onto the EVOTIP (EV-2013 EVOSEP) following the manufacturer’s instructions. The analytical column was a 15cm x 150μm ID, with Dr Maisch C18 AQ, 1.5 μm beads (EV-1137 EVOSEP) with a column oven set at 40°C. The eluents were 0.1% formic acid in water and 0.1% FA in ACN. The Evosep One method was 15 SPD (88 min gradient)^74^, and the flow rate 0.22 μl/min. The mass spectrometer was operated in data independent acquisition mode (DIA). Full MS1 scans were acquired in the Orbitrap with a scan range of 350 - 1200 m/z and a resolution of 120,000 (at 200 m/z). Automatic gain control (AGC) was set to a target of 1 x 106 and a maximum injection time of 56 ms. MS2 spectra were acquired in DIA mode using 33 m/z variable windows with 0.5 m/z overlap. MS2 spectra were analyzed in the Orbitrap with a resolution of 30,000 (at 200 m/z). A higher energy collision induced dissociation (HCD) method (28% NCE) was applied with an AGC target of 5x104 and a maximum injection time of 55 ms. Orbitrap Eclipse Tune Application 4.2.4321 and Xcalibur version 4.7.69.37 were used to operate the instrument and to acquire data. The mass spectrometry proteomics data has been deposited to the ProteomeXchange Consortium via PRIDE^75^. The raw files obtained were analyzed using Data-Independent Acquisition by Neural Networks software (DIA-NN, v1.9.2 beta 27)^76^, searching against the reviewed SwissProt database for human (retrieved on January 2025), miniTurbo-v5-SS18::SSX2 constructs and common contaminants (retrieved on February 2023) in library-free mode. Acetylation in protein N-terminal, oxidation in Methionine and Methionine excision at protein N-terminal were accepted as variable modifications, none modifications were set up as fixed modification. The maximum number of tryptic missed cleavages and the maximum number of variable modifications allowed were set to 2, respectively. The m/z range for the precursors and the fragment ions were set to 350 - 1200 and 200 - 2000, respectively. The match between runs option and the heuristic algorithm for the protein inference at the isoform IDs level option were also enabled. The remaining DIA-NN parameters were kept at their default values. The resulting search output was preprocessed, visualized and statistically analyzed using the R programming language (https://www.r-project.org/). The package diann-rpackage (https://github.com/vdemichev/diann-rpackage) was used to load DIA-NN search report and to generate the protein groups MaxLFQ quantification^77^ prior filtering by precursor Q value and protein group Q value below 0.01. Then, contaminant protein groups and protein groups quantified with less than two peptides were filtered out. Additionally, no missing values in at least one of the conditions in each contrast (SSX2 and mT), Then, MaxLFQ were log2-transformed and the remaining missing values were imputed with normally distributed random numbers with *μ*imputed = *μ* - 1.8 *σ* and *σ*imputed = 0.3 *σ*, where *μ* and *σ* are the non-missing values mean and standard deviation, respectively. The subsequent differential protein abundance analysis was performed leveraging the “limma” package^72^. Contrast SSX2 vs mT was defined and cutoffs for the fold change and adjusted p-value (*p*adj < 0.05) were defined to establish significant proteins.

### *Smarca4* deleted mouse models

All mouse experiments were conducted under the approval of the University of Utah Institutional Animal Care and Use Committee. The *Smarca4-floxed* mouse strain was obtained from Terry Magnuson^78^. The *hSS1* and *hSS2* mouse strains were generated in the laboratory^9^. Experimental tumorigenesis mice were bred from parents that were homozygous for either *hSS1* or *hSS2* and heterozygous for Smarca4-floxed, such that littermates included homozygous wildtype, heterozygous, and homozygous floxed alleles in 1:2:1 Mendelian ratios. Mice were injected in the hindlimb with TATCre protein at age 8 days, then monitored for tumorigenesis by inspection and later caliper measurements at least twice each week. After tumors reached morbid size, mice were euthanized and tissues were harvested by necropsy. Tissue blocks were fixed in formalin and paraffin embedded prior to sectioning and staining with hematoxylin and eosin. Review of H&E stained sections was performed under light microscopy by one blinded to genotype of the previous host mouse.

### SS18 AlphaFold3 interaction screen

Paired and unpaired multiple sequence alignments for UniProt canonical sequences of proteins present across IP/MS datasets (N=2,186) were generated with LocalColabFold, running ColabFold version 1.5.5 on default settings^79^. A local version of AlphaFold3^80^ was run on a single RTX 6000 Ada GPU (CUDA version 12.6).

### Immunofluorescence staining

0.3 × 10^6^ HEK293T cells were seeded 24h before transduction with pLV-EF1α-eGFP constructs in 6-well plates containing coverslips. Cells were transduced and treated with 500nM ACBI1, 500nM dCBP-1 or DMSO for 72 h and were then fixed with 4% paraformaldehyde for 10 min. Permeabilization was performed using Triton X (0.1% in PBS) for 12 min, followed by incubation with blocking solution (1% BSA, 0.1% gelatin fish in PBS) for 1 h. Incubation with primary antibodies was performed in blocking buffer at room temperature (20–22°C) for 1 h. Cells were incubated with secondary antibodies for 1 h at room temperature. Cells were then mounted in VECTASHIELD Antifade Mounting Medium containing 4ʹ,6-diamidino-2-phenylindole (DAPI; Vector Laboratories).

### Image capture, processing and analysis

Confocal images were acquired on a Leica TCS SP5 inverted confocal microscope using an HCX PL APO 63x/1.40-0.60 Oil Lbd BL objective, and a single z-stack was captured. Samples were imaged using 405, 488, 561, 594 and 633 nm laser lines using sequential mode in the Leica Application Suite software at 12bit. Samples were imaged using a 512×512 format at a speed of 100 Hz using line averaging at 4 with a zoom factor of 5 for showing three or four nuclei. Images were then smoothed and adjusted for brightness and contrast using the ImageJ/Fiji software. Between 10 and 30 foci were analyzed by drawing a line of 5um centered on the H2Aub-rich foci or eGFP-rich foci. Using the “plot profile” tool on ImageJ/Fiji, the fluorescence intensity across that line was measured for the two chanels of interests: 488 (eGFP) and 594 (H2Aub, EP300 or SMARCC1). The data of the 2 replicates was combined and plotted using GraphPad prism, which was also used to computed the Pearson correlation between the channels.

### Statistics and reproducibility

Details of the individual statistical analyses and tests, as well as the number of biological replicates, can be found in the respective figure legends. Statistical analysis was performed using Perseus, GraphPad Prism and R.

### Data availability

HSSY-II ChIP sequencing data reanalyzed in Fig. 2 originates from GEO accession number GSE108929. KHOS-240S CUT&RUN reanalyzed in Fig3 originates GEO accession number GSE205955.

## AUTHORS’ CONTRIBUTIONS

A.S. designed, performed, and analyzed the experiments, generated figures and wrote the manuscript. S.M and L.M performed and analyzed experiments. E.C.W., S.B. assisted with experiments. J.L., K.S-F., L.M., L.C., B.R.C. and K.B.J. designed, performed and analyzed the *Smarca4* deletion sarcomagenesis data. J.A.K., S.T. and C.M-R. designed, performed and analyzed the miniTurbo IP-MS data. A.K.J. provided materials and information for IP-MS experiment in ACBI1 treated HEK293T cells expressing SS18::SSX1 and assisted with experiments. M.S. and D.F. analyzed the mass spectrometry data. M.B., J.A.M. performed and provided the Alpha Fold 3 analyses. N.S.B., conceived and co-ordinated the study; designed, performed, and analyzed experiments; acquired funding and wrote the manuscript A.B. conceived and coordinated the study; designed experiments; acquired funding and wrote the manuscript. All authors read, revised and approved the final manuscript for publication.

## ACKNOWLEDGEMENTS

We thank the Flow Cytometry, Genomics and Proteomics core facility of DKFZ, the Advanced Imaging Resource facility of the Institute of Genetics and Cancer and the MS & Proteomics facility of IRB Barcelona for their exceptional support and service. We express our gratitude to Ralph S. Grand and Martina Capriati (ZMBH, Heidelberg) for their help with the ATAC-sequencing protocol and to the lab of Priya Chudasama and Stefan Fröhling for providing the CME-1 synovial sarcoma cell line. We acknowledge funding from the CGC PROTECT. A.S. is supported by the DKFZ International PhD Program. J.A.W. is supported by the EU’s Horizon Europe programme (MSCA Postdoctoral Fellowship 2023 -101154879). Work was also supported by the L. B. and Olive S. Young Presidential Endowed Chair for Cancer Research, the Huntsman Cancer Foundation, and the National Cancer Institute of the National Institutes of Health under award numbers U54CA231652 and P30CA042014. E.C.W. is supported by the IGC Langmuir Talent Development Fellowships in Cancer Research and N.S.B is supported by a University of Edinburgh Chancellor’s Fellowship. Work in the laboratory of N.S.B. was supported by the Bone Cancer Research Trust (BCRT10124) and SarcomaUK (SUK13.2024). This project received funding from the European Research Council (ERC) under the European Union’s Horizon 2020 research and innovation program (grant agreement n° 805338 [A.B.] and 101001169 [J.A.M.]).

---

**Figure S1.**
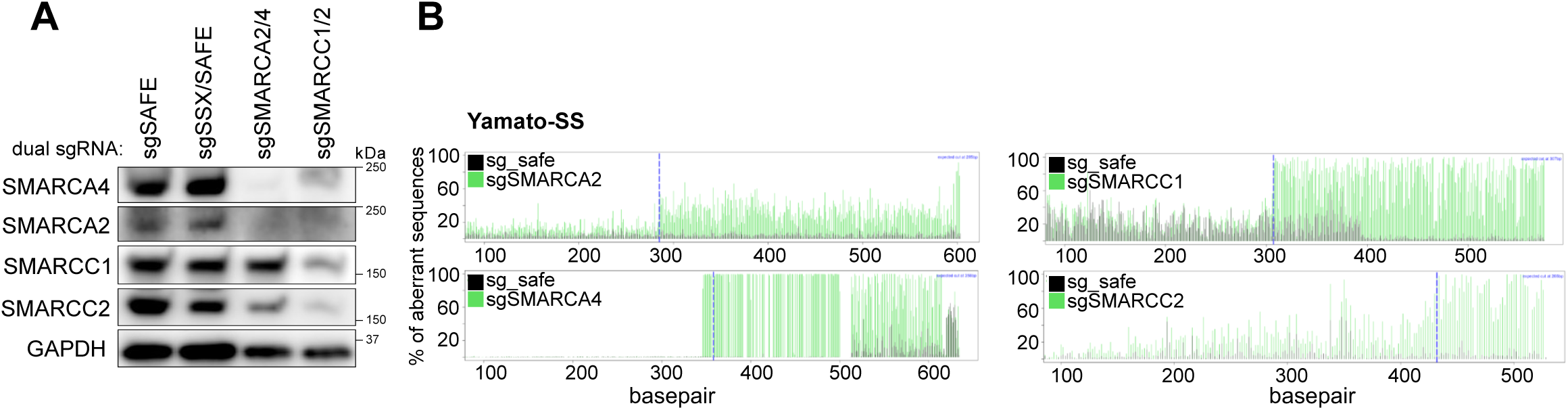
**(A)** Western Blot of KHOS-240S-Cas9 whole cell extracts expressing a safe sgRNA as control or with double guides targeting SSX + a control region (sgSSX/SAFE), SMARCA2 + SMARCA4 (sgSMARCA2/4) or SMARCC1 + SMARCC2 (sgSMARCC1/2) revealed using SMARCA4, SMARCA2, SMARCC1, SMARCC2 or GAPDH antibodies. **(B)** Tracking of Indels by Decomposition (TIDE) assay displaying the percentage of aberrant sequences after Cas9 editing for guides targeting SMARCA2, SMARCA4, SMARCC1 and SMARCC2 versus the wild-type sequence (sgSAFE guide) in Yamato-SS cell line.

**Figure S2.**
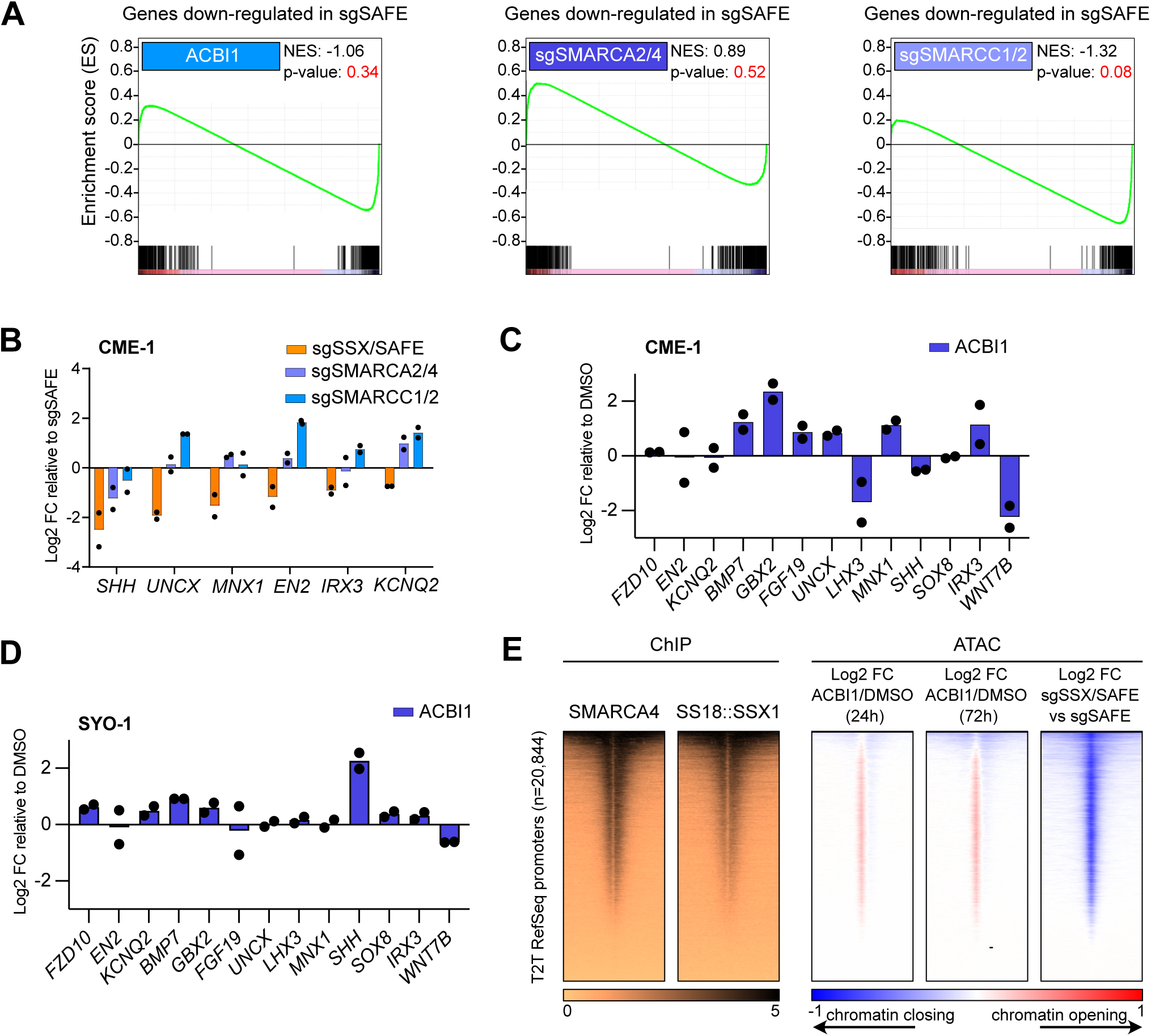
**(A)** Gene set enrichment analysis (GSEA) comparing differentially expressed genes in the ACBI1, sgSMARCA2/4 or sgSMARCC1/2 RNAseq data in HSSY-II to the top 500 genes downregulated in sgSSX. **(B)** qRT-PCR displaying log2-transformed fold change of mRNA levels normalised by GAPDH in CME-1-Cas9 cells expressing sgSAFE, (sgSSX/SAFE), (sgSMARCA2/4) or (sgSMARCC1/2) for 7 days relative to sgSAFE. Data represents the mean of n = 2 biological replicates. **(C)(D)** qRT-PCR displaying log2-transformed fold change of mRNA levels normalised by GAPDH in CME-1 (C) or SYO-1 (D) cells treated for 72h with DMSO or 500nM ACBI1 relative to DMSO. Data represents the mean of n = 2 biological replicates. **(E)** Left: heatmaps for SMARCA4 and SS18::SSX1 (endogenously HA tagged) ChIP-seq from Banito et al., 2018 over T2T hs1 Refseq promoters (n = 20,844). Rows correspond to ±5-kb regions across the midpoint of each SMARCA4-enriched region, ranked by increasing signal. Right: ATAC-seq heatmaps displaying log2-transformed fold change of ACBI1 treated cells over DMSO after 24h or 72h of treatment and of sgSSX/SAFE expressing cells over sgSAFE cells.

**Figure S3.**
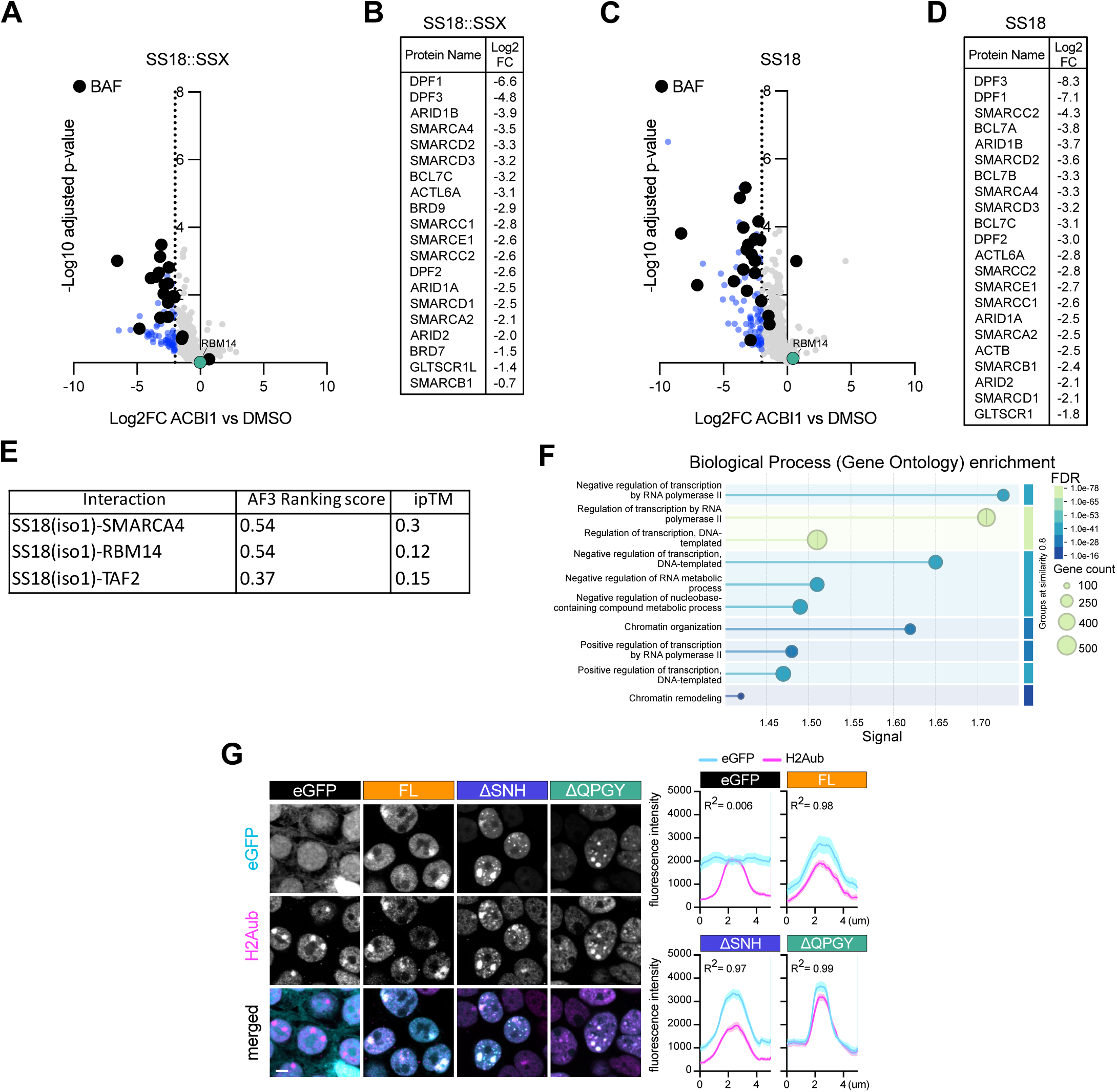
**(A)(C)** Volcano plots of mass spectrometry data following eGFP-SS18::SSX (A) or eGFP-SS18 (C) pull down in ACBI1 treated HEK293T cells normalised to interactome from DMSO treated cells and to eGFP-SSX pull down. Data represents log2 fold change enrichment plotted on the x-axis and -log10 (adjusted p value) plotted on the y-axis. Blue dots indicate lost interactors upon ACBI1 treatment, Black dots indicate BAF members. n = 3 biological replicates. **(B)(D)** List of BAF members present in interactome dataset and their associated log2 fold change value. Listed members are shown as black dots in (A) and (C). **(E)** Alphafold predictions scores of SS18 (iso1) with SMARCA4, RBM14 and TAF2. ipTM (interface predicted TM-score) is displayed. **(F)** Gene ontology enrichment analysis of the significantly enriched proteins from the mT-v5-SS18::SSX2 proteomics in HEK293T cells. Analysis was conducted using STRING (v12). The list of total proteins detected in proteomics was used as the statistical background for the enrichment analysis. p-value < 0.05 and log2 fold change > 1 set as significantly enriched. **(G)** Left: Immunofluorescence of HEK293T cells expressing eGFP, FL, ΔSNH or ΔQPGY (cyan) stained for H2AK119ub1 (H2Aub, magenta). Images are representative of n = 2 biological replicates. Scale bar, 5 μm Right: Profile of fluorescence intensity of eGFP, FL, ΔSNH or ΔQPGY over H2Aub foci. Data represent the mean ± S.E.M of n = 2 biological replicates. For each replicate and condition, > 10 foci where profiled. R² indicates Pearson correlation coefficient.

**Figure S4.**
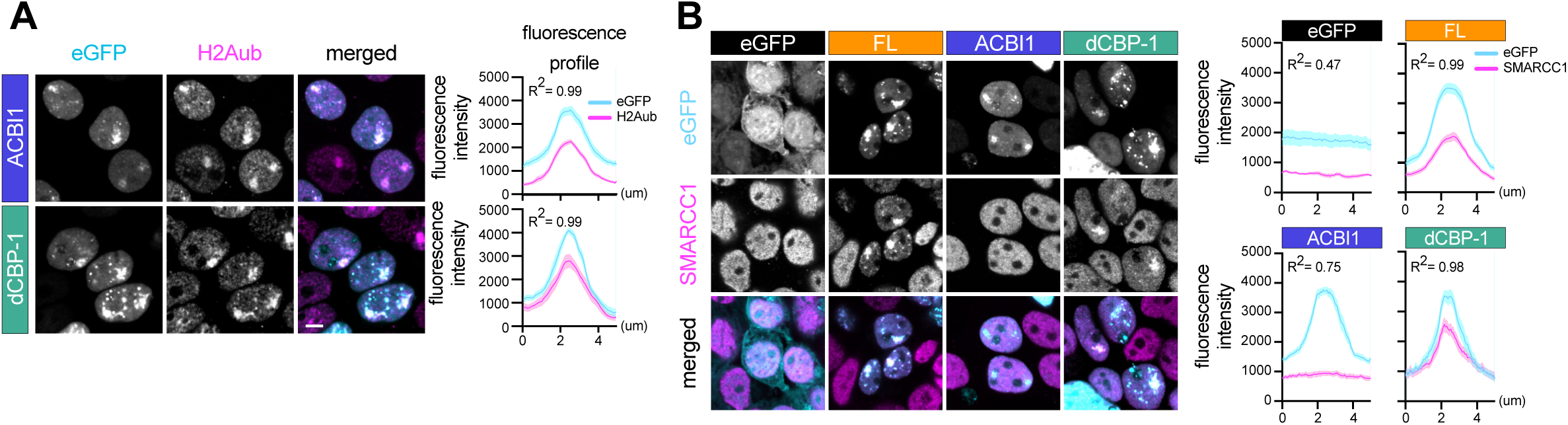
**(A)** Left: Immunofluorescence of HEK293T cells expressing wild-type SS18::SSX1 (FL) treated for 72h with either 500nM ACBI1 or 500nM dCBP- 1 (cyan) stained for H2AK119ub1 (H2Aub) (magenta). Images are representative of n = 2 biological replicates. Scale bar, 5 μm. Right: profile of fluorescence intensity of FL, treated for 72h with either 500nM ACBI1 or 500nM dCBP-1 over H2Aub or eGFP foci. Data represent the mean ± S.E.M of n = 2 biological replicates. For each replicate and condition, > 10 foci where profiled. R² indicates Pearson correlation coefficient. **(B)** Left: Immunofluorescence of HEK293T cells expressing eGFP or wild-type SS18::SSX1, either untreated (FL) (cyan) or treated for 72h with 500nM ACBI1 or 500nM dCBP-1 stained for SMARCC1 (magenta). Images are representative of n = 2 biological replicates. Scale bar, 5 μm. Right: Profile of fluorescence intensity. Data represent the mean ± S.E.M of n = 2 biological replicates. For each replicate, > 10 foci where profiled. R² indicates Pearson correlation coefficient.

